# Splenic follicular B cells promote adverse cardiac remodeling after myocardial infarction via MHC II-dependent antigen presentation

**DOI:** 10.1101/2024.05.08.593153

**Authors:** Obialunanma V. Ebenebe, Jana P. Lovell, Carolina Duque, Yiran Song, Sylvie Rousseau, Kshipra Keole, Sogol Sedighi, Aashik Bhalodia, Kevin Bermea, Charles D. Cohen, Monica R. Mugnier, D. Brian Foster, Scott Krummey, Simone Sidoli, Luigi Adamo

**Affiliations:** Division of Cardiology, Department of Medicine, Johns Hopkins University School of Medicine, Baltimore, Maryland; W. Harry Feinstone Department of Molecular Microbiology and Immunology, Johns Hopkins Bloomberg School of Public Health; Department of Pathology, Johns Hopkins University School of Medicine, Baltimore, Maryland; Department of Biochemistry, Albert Einstein College of Medicine, Bronx, New York

**Author notes:** Address of Correspondence: Luigi Adamo, MD, PhD, Division of Cardiology, Johns Hopkins University School of Medicine 720 Rutland Avenue, Ross Research Building 809 Baltimore, MD 21205, @luigiadamomdphd. these two authors share first authorship (Obialunanma V. Ebenebe, Jana P. Lovell). **Sources of Funding:** This study has been funded through NHLBI grants 5K08HLO145108-03 and 1R01HL160716-01 to L.A. and T32-HL007227 to O.V.E and J.P.L and institutional funds from the Johns Hopkins Division of Cardiology awarded to L.A. **Disclosures:** L.A. is co-founder of i-Cordis, LLC, a startup company focused on the development of immunomodulatory therapies for heart failure, and he is a consultant for Novo Nordisk and Kiniksa Pharmaceuticals.

## Abstract

The cardio-splenic axis is a promising therapeutic target in ischemic heart failure (HF), but splenic–cardiac interactions after myocardial infarction (MI) are poorly understood. Here, we show that splenic follicular B cells drive ischemic HF via MHC class II–mediated antigen presentation. Transfer of splenic B cells from mice with ischemic HF induced adverse cardiac remodeling and modulated myocardial T-cells in naïve recipients. Single-cell RNA sequencing of mouse and human post-MI B cells revealed upregulation of antigen-presentation related pathways. Mass spectrometry of the MHC II peptidome demonstrated enrichment of cardiac-derived peptides in MHC II molecules of splenic B cells from ischemic HF mice. B cell-specific deletion of MHC II suppressed adverse cardiac remodeling after adoptive transfer of post-MI splenic B cells. These results broaden our understanding of B-lymphocyte biology and point towards MHC II-mediated signaling in B-cells as a novel therapeutic target in ischemic HF.

---

Acute myocardial infarction (MI) triggers a robust activation of inflammatory responses [1, 2]. These inflammatory responses are physiological and necessary to limit tissue injury and induce tissue repair [3, 4]. However, sustained inflammation contributes to adverse cardiac remodeling and thus, to the development and progression of ischemic heart failure [5]. Despite advancements in our understanding of the relationship between chronic inflammation and heart failure (HF), clinical trials investigating immunomodulatory therapies in HF have yielded largely unsuccessful results. The failure of these therapies is partly due to the inability to selectively target maladaptive rather than protective inflammatory responses post-MI [6].

As such, an improved understanding of the sustained maladaptive inflammatory responses contributing to the pathogenesis of chronic HF is needed. Prior murine and human data indicate that acute MI induces splenic remodeling and mobilization of immune cells from the spleen to the injured myocardium [7]. Activation of this “cardio-splenic axis” has been shown to contribute to adverse cardiac remodeling and the development of chronic HF [8]. Thus, the cardio-splenic axis has been identified as a promising therapeutic target for developing immunomodulatory treatments for HF. However, our mechanistic understanding of the processes by which specific splenic immune cells contribute to adverse cardiac remodeling remains limited.

B cells comprise the majority of immune cells within the spleen of most vertebrates. B cells are also prevalent in healthy hearts and can recirculate between the spleen and the heart [9–11]. Growing evidence shows that B cells are intricately involved with cardiac function and dysfunction, and following acute MI, contribute to the development and progression of ischemic HF [12–16]. However, to date, the precise role of B cells within the cardio-splenic axis in HF has not been systematically investigated.

Here, we show that B cells, specifically splenic follicular B cells, play an essential role within the cardio-splenic axis of ischemic heart failure by promoting adverse cardiac remodeling post-MI via MHC II-dependent antigen presentation.

## Results

### Splenic B cells contribute to adverse cardiac remodeling in ischemic heart failure

To investigate whether splenic B cells are involved in the adverse cardiac remodeling of ischemic HF, we designed adoptive transfer experiments. Unfractionated splenocytes or purified splenic B cells were transferred from HF mice into naïve mice via intravenous injection. Splenic immune cells isolated from sham-operated animals were used as controls. The naïve recipient mice were monitored for eight weeks after adoptive transfer (Fig.1a).

**Fig. 1:**
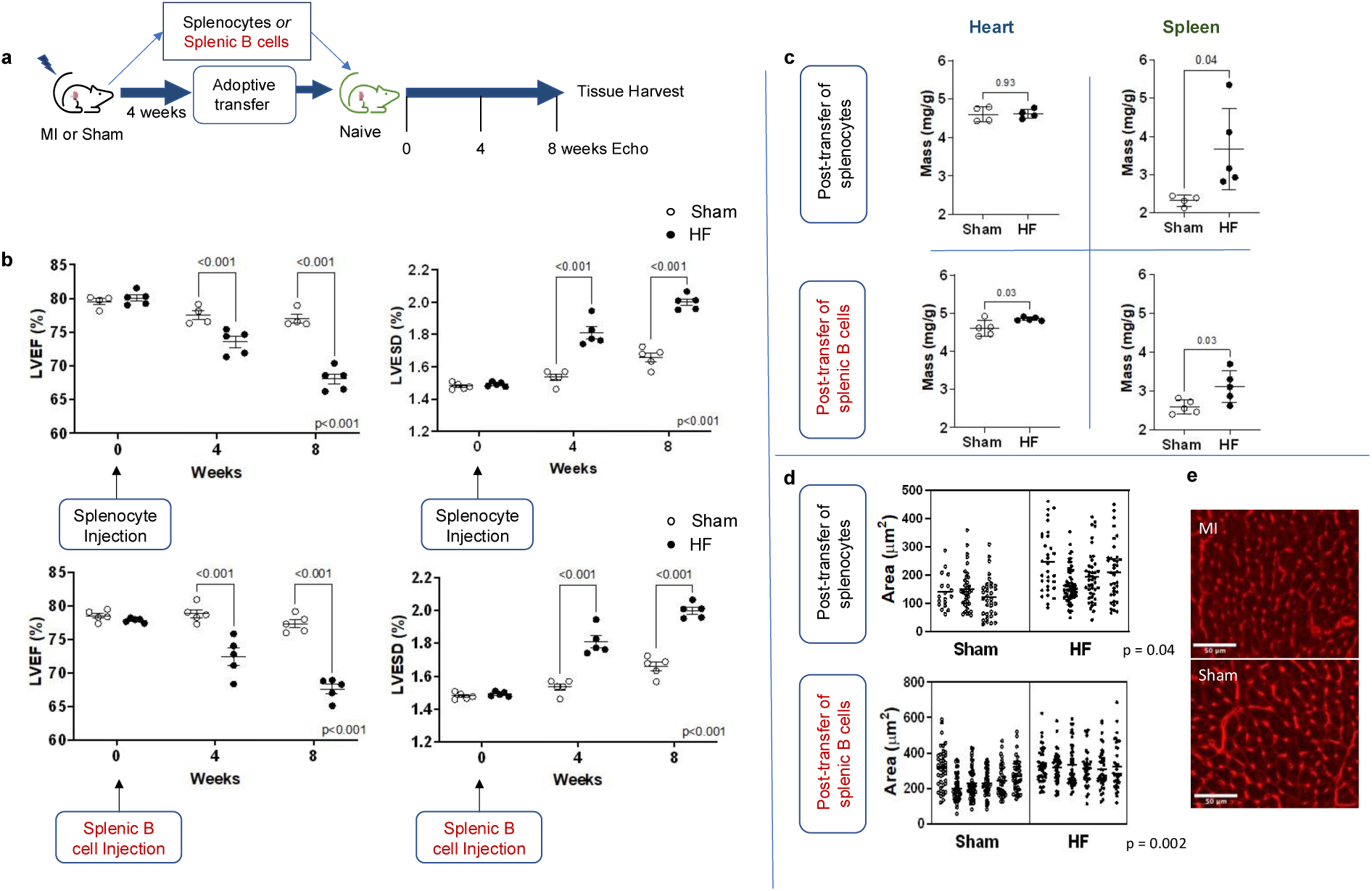
Splenic B cells promote adverse cardiac remodeling in ischemic heart failure. **a**) Schema for adoptive transfer studies. Four weeks following permanent LAD coronary ligation or sham surgery, unfractionated splenocytes or isolated splenic B cells were transferred to naïve recipient mice. Recipient mice underwent serial transthoracic echocardiography over 8 weeks followed by tissue harvest. **b**) On echocardiography, adoptive transfer of splenocytes or isolated splenic B cells from heart failure (HF) mice resulted in significant reduction in left ventricular ejection fraction (LVEF) and increase in left ventricular end-systolic diameter (LVESD) in naïve recipient mice. **c**) Quantitative gravimetric data revealed significantly larger hearts and spleens (normalized to body weight) in naive mice 8 weeks post-adoptive transfer of splenocytes or isolated splenic B cells from HF mice relative to recipients of splenocytes or splenic B cells from sham-operated mice. **d**) Cardiomyocyte area was increased in recipients of splenocytes or isolated splenic B cells from HF mice relative to sham recipients; n = 5-6 mice per group. Mean values ± SD are represented. **e**) Representative images of WGA staining of hearts from recipients of isolated splenic B cells from HF or sham-operated mice. LAD- left anterior descending. HF-heart failure.

Consistent with previously published data [8], we found that adoptive transfer of splenocytes from mice with ischemic HF resulted in adverse cardiac remodeling in recipient mice. From serial echocardiography, recipients of splenocytes from mice with HF had reduced left ventricular ejection fraction (LVEF) and increased left ventricular end-systolic diameter (LVESD) eight weeks post adoptive transfer compared to recipients of splenocytes from sham-operated mice (Fig.1b; mean LVEF 68.1±1.7% vs 77.0±1.3% HF vs. sham recipient, q<0.001 at 8 weeks; mean LVESD 2.01±0.7mm vs. 1.67±0.02mm HF vs. sham recipient, q<0.001 at 8 weeks).

To investigate the specific role of splenic B cells within the cardio-splenic axis following ischemic myocardial injury, we performed adoptive transfer of purified splenic B cells from HF or sham-operated mice (≈95% isolated B cell purity, Supplementary Fig.1). Over the 8-week period, the adoptive transfer of isolated splenic B cells from mice with HF produced a degree of adverse remodeling similar to that observed with the adoptive transfer of unfractionated splenocytes (Fig.1b, bottom panels). Recipients of splenic B cells from mice with HF experienced a progressive reduction in LVEF and progressive LV dilatation, detected at 4- and 8-weeks post adoptive transfer (Fig.1b; mean LVEF 67.6±1.6% vs. 77.3±1.3% HF vs. sham recipient, q<0.001 at 8 weeks; mean LVESD 2.00±0.05mm vs. 1.66±0.06mm HF vs. sham recipient, q<0.001 at 8 weeks). Further experiments wherein we extended the post adoptive transfer monitoring revealed that this effect was progressive to 16-weeks post adoptive transfer (Supplementary Fig.2).

Adverse cardiac remodeling in naïve recipients of splenocytes and splenic B cells isolated from HF mice was further indicated by gravimetric analyses of the heart and spleen, as well as wheat-germ agglutinin (WGA) staining of myocardial cross-sections. Gravimetric analysis revealed increased splenic and cardiac mass in recipients of splenic B cells from mice with HF compared to recipients of B cells from sham-operated mice 8 weeks following adoptive transfer (Fig.1c; for purified splenic B cells: spleen mean 3.11±0.4 mg/g vs 2.59±0.2 mg/g, p = 0.03; heart mean 4.85±0.05 mg/g vs 4.60±0.2 mg/g, p = 0.03; Supplementary Fig.3 displays the combined gravimetric data from multiple independent experiments). Furthermore, WGA staining of myocardial cross-sections showed that increased heart mass was associated with cardiomyocyte hypertrophy (Fig.1d; p = 0.002).

### Adoptively transferred splenic B cells are distributed between the spleen and heart

To investigate the dynamics of adoptively transferred splenic B cells, we repeated the adoptive transfer experiments using animals carrying CD45.1 isoform as donors and animals carrying CD45.2 isoform as recipients (Fig.2a). Donor CD45.1 B cells were present in the peripheral blood of naïve recipient mice at 1- week and 8- weeks following adoptive transfer (Fig.2b; ∼3% and ∼2% of circulating donor-derived B cells at 1- and 8- weeks, respectively). Flow cytometry of the spleen and heart 8 weeks post-adoptive transfer also revealed the presence of donor CD45.1 B cells, indicating successful engraftment of CD45.1 B cells in recipient mice (Fig.2b; Supplementary Fig.4 also includes non-adoptive transfer controls).

**Fig. 2:**
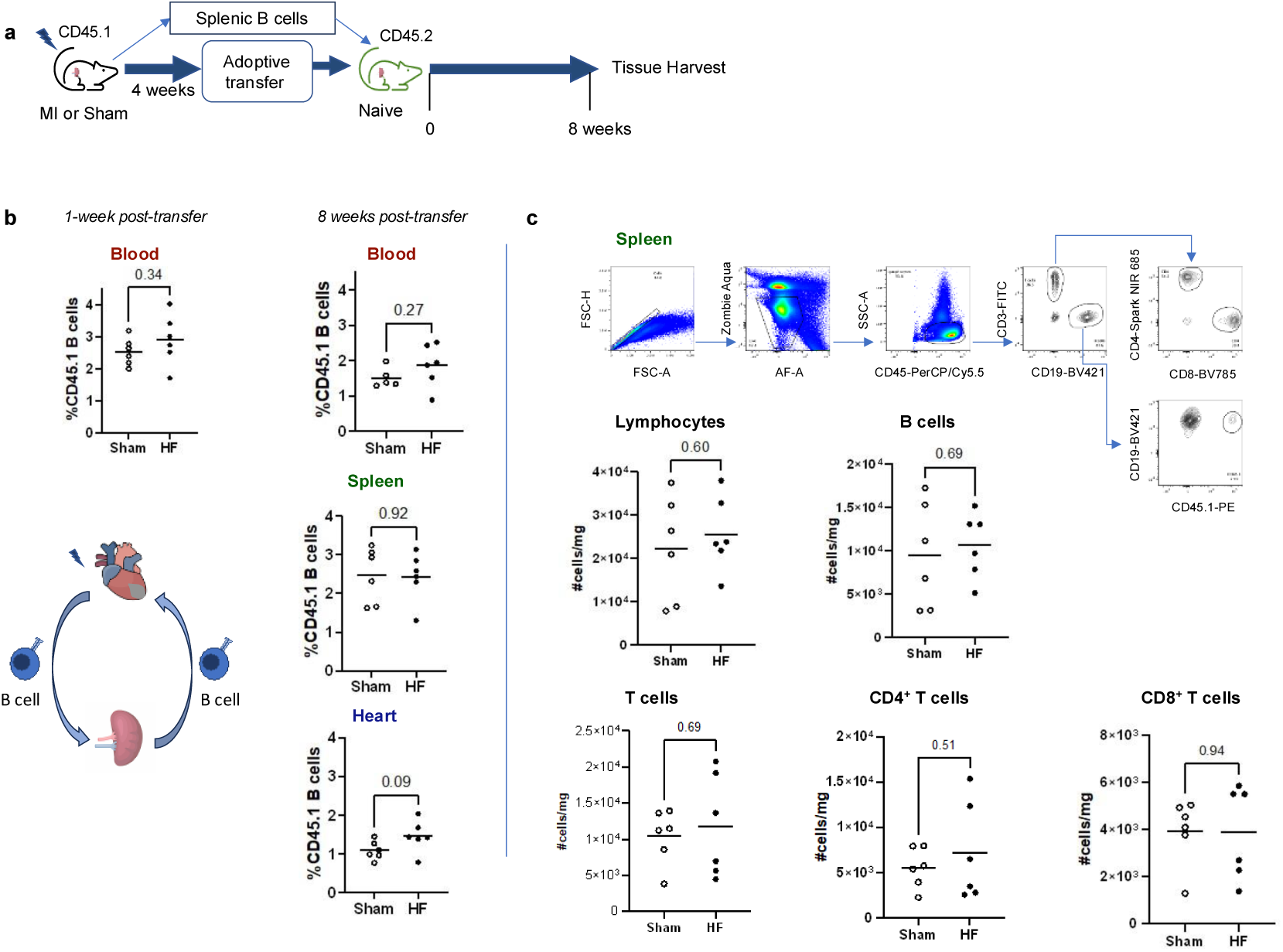
Adoptively transferred splenic B cells from mice with ischemic HF are found in the heart and spleen of recipients, and do not alter the immune composition of the spleen of naïve mice. **a**) Schema for adoptive transfer of isolated splenic CD45.1 B cells 4 weeks following LAD ligation or sham surgery to naïve CD45.2 mice. **b**) Flow cytometry of peripheral blood from naïve recipient mice revealed the presence of donor CD45.1 B cells 1 week and 8 weeks following adoptive transfer. Donor CD45.1 B cells were also present in the spleen and heart of CD45.2 mice 8 weeks after adoptive transfer. **c**) Representative gating strategy for flow cytometry of spleens from recipient mice 8 weeks post-adoptive transfer of splenic B cells. Adoptive transfer of splenic B cells from HF mice had no effect on total number of lymphocytes, B cells, CD4^+^ and CD8^+^ T cells relative to recipients of sham splenic B cells. LAD- left anterior descending. HF-heart failure. n = 6 mice per group. Mean values are represented.

### Ischemic myocardial injury results in differential expression of antigen processing and presentation pathways in splenic B cells

To explore the mechanistic relationship between the adoptive transfer of splenic B cells from mice with ischemic HF and adverse cardiac remodeling in naïve mice, we first analyzed the recipients’ spleen and heart via high-dimensional flow cytometry. At the splenic level, adoptive transfer of splenic B cells from mice with ischemic HF did not affect the numbers of splenic lymphocytes or the prevalence of B cell subsets (Fig.2c, Extended data Figure 1); nor were there changes in splenic CD3 T cells, CD4 T cells, and CD8 T cells numbers relative to recipients from sham-operated mice (Fig.2c). Closer examination of splenic T cell subsets indicated minor differences. We noted a downward trend in the percentage of memory CD4 T cells expressing CTLA-4 and FOXP3 (CD44^hi^, CD62L^lo^, CTLA-4^+^, FOXP3^+^) and a corresponding increase in memory CD4 T cells negative for CTLA-4 and FOXP3 markers (CD44^hi^, CD62L^lo^, CTLA-4^-^, FOXP3^-^, Extended Data Fig.2). We did not detect differences in subsets of splenic CD8 T cells (Extended Data Fig.3). At the myocardial level, via flow cytometry, we did not detect differences in cardiac immune cell populations between recipients of splenic B cells from mice with ischemic HF and recipients of splenic B cells from sham-operated mice (Extended Data Fig.4).

Since this analysis was unrevealing, we focused our attention on the donors; the population of splenic B cells that were being adoptively transferred. We first used high-dimensional flow cytometry. Within the spleen, B cells can be sub-classified based on specific cell markers [17, 18]. There were no changes in the percentage of splenic B cell subtypes isolated from mice with ischemic HF (4 weeks post-MI surgery) compared to sham-operated mice (Extended Data Fig.5). We then used single-cell RNA sequencing (scRNAseq) to more deeply probe the molecular character of the donor splenic B cells from HF mice (Fig.3a). V(D)J analysis did not reveal an expansion of specific splenic B cell clonotypes in ischemic HF (4 weeks post-MI). Suggesting that the effects observed at the time of adoptive transfer were not due to splenic B cells’ clonal expansion (Fig.3b; Supplementary Fig.5). UMAP plots were then used to visualize the clustering of gene expression profiles of splenic B cells (Fig.3c). Comparative gene expression analysis revealed several genes differentially expressed in splenic B cells from mice with ischemic HF (4 weeks post-MI surgery) (Supplementary Table 1).

**Fig. 3:**
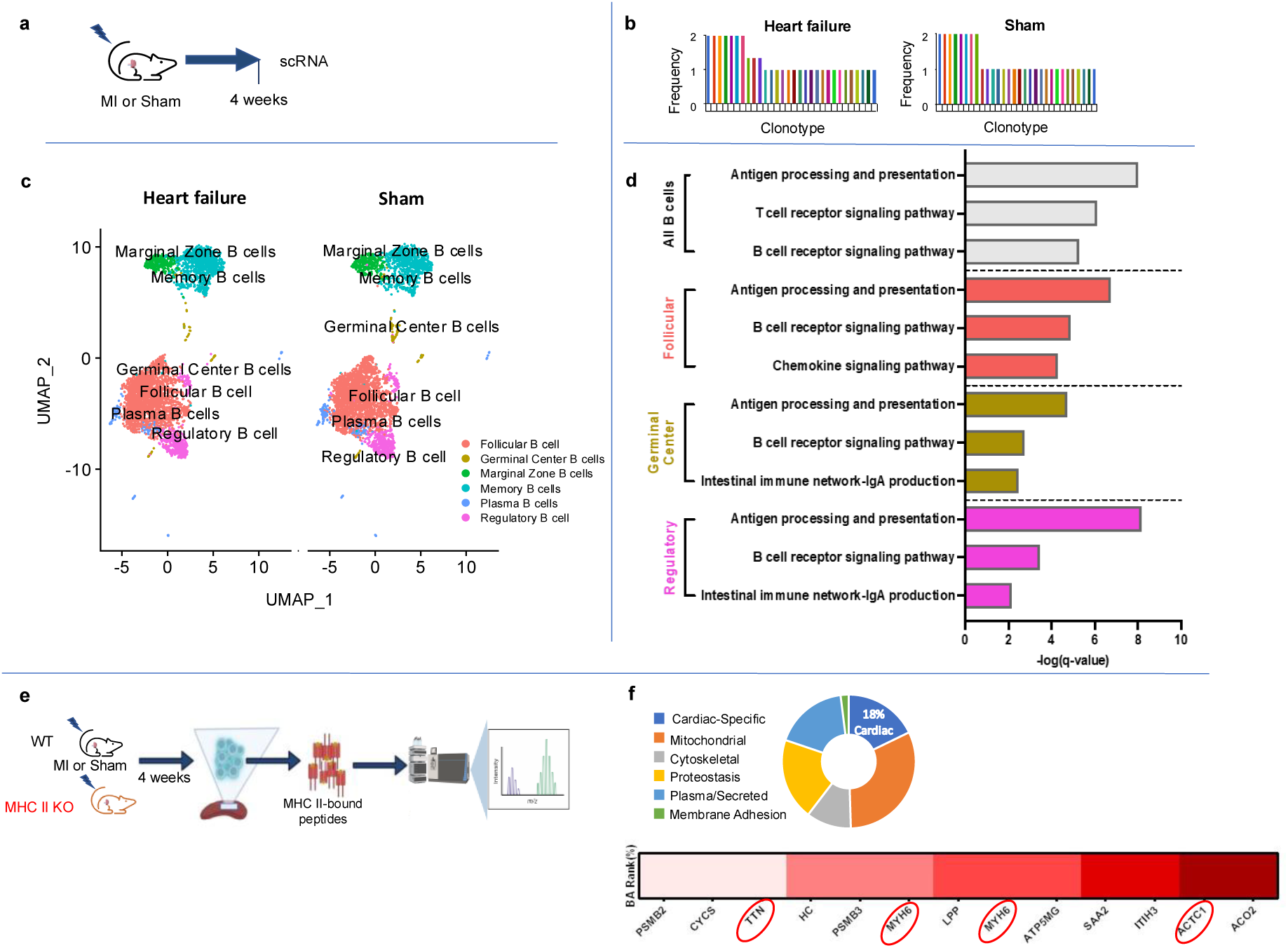
Ischemic myocardial injury is associated with activation of antigen processing and presentation pathways in splenic B cells and with enrichment of myocardial peptides within the MHC-II molecules of these B cells. **a**) Schema for single cell sequencing of splenic B cells from mice 4 weeks following permanent LAD coronary ligation or sham surgery. **b**) V(D)J analysis of splenic B cells. Ischemic myocardial injury did not induce splenic B cell clonal expansion. **c**) UMAP plots of splenic B cells from heart failure or sham surgery mice. Splenic B cells were classified into B cell sub-types. **d**) KEGG pathway analysis of differentially expressed genes in splenic B cells (fold change >1, p<0.05) revealed significant differential expression of antigen processing and presentation in multiple B cell sub-types, post-MI. All pathways with FDR <0.05 are listed. (n = 4-5 mice per group). **e**) Schema for mass spectrometry of MHC-II bound peptides from splenic B cells four weeks after permanent LAD coronary ligation or sham surgery. Splenic B cells from MHC II KO (*Cd19tm1(cre)Cgn/-H2-Ab1b-*tm1Koni/ *b-*tm1Koni) mice were included besides cells from WT (wildtype) mice to allow identification of background signal. **f**) Categories of proteins both enriched > 1.2 folds in the MHC-II molecules of splenic B cells isolated from mice with ischemic HF and *in silico* confirmed binding to the C57Bl/6 specific MHC-II molecule H2-I-Ab. 18% of these proteins are cardiac-specific. The peptides/corresponding proteins identified as strong binders are reported in the heat map. The cardiac proteins titin (TTN), myosin heavy chain 6 (MYH6) and α-actin (ACTC1) were identified among the top predicted strong binding peptides. MYH-6 is repeated twice because two different peptides from MYH-6 were detected. LAD- left anterior descending. HF-heart failure.

To evaluate the biological relevance of these differentially expressed genes (DEGs), we performed pathway analyses using KEGG pathway annotation [19, 20]. Antigen processing and presentation was the top biological pathway identified as differentially regulated (q-value = 8.02E-9; Fig.3d). Some of the specific DEGS included HSPA8, HSP90AA1, HSP90AB1, which encode heat shock proteins (HSPs) involved in MHC-II mediated antigen presentation [21, 22]; H2-Eb1 and H2-DMb1 which encode MHC-II proteins and IFI30 (also known as GILT) which is involved in MHC-II mediated antigen presentation [23]. Further evaluation of B cell subtypes revealed significant differential regulation of the antigen processing and presentation pathway in the follicular, germinal center, regulatory, plasma, and memory B cells (Fig.3d) but not in marginal zone B cells (Supplementary Table 2).

To further corroborate these findings, we expanded our analysis to myocardial B cells and peripheral blood B cells in animals with ischemic HF (Supplementary Figs.6 and 7). We also included animals with recent MI (four days post-MI surgery) to gain longitudinal insight (Supplementary Figs.8-10). These analyses also pointed towards antigen processing and presentation as one of the top sustained, differentially regulated pathways in myocardial and circulating B cells in response to myocardial injury. More specifically, antigen processing and presentation was differentially regulated in multiple cardiac and peripheral blood, B cell subtypes from mice with ischemic HF (Supplementary Figs.6 and 7), with an especially strong signal in myocardial and circulating follicular B cells (Supplementary Figs.6C and 7C). Recent MI was compared to the 8-weeks post sham surgery (which is a suboptimal control); yet this analysis also highlighted significant differential regulation of antigen processing and presentation in splenic, cardiac, and peripheral blood B cells with a predilection for follicular B cells (Supplementary Figs.8-10). Lists of the DEGs associated with antigen processing and presentation in mice with ischemic HF and mice with recent MI are provided within the spreadsheets included in Supplementary Table 1.

### MHC II molecules of splenic B cells from mice with ischemic HF are loaded with cardiac peptides

Collectively, our transcriptomic data suggested that splenic B cells might modulate adverse cardiac remodeling in ischemic HF via major histocompatibility complex class II (MHC II) mediated antigen presentation. This was an unexpected finding. B cells have three main functions: antibody production, cytokine secretion, and antigen presentation. It has been reported that B cells may contribute to adverse cardiac remodeling following MI via the production of autoantibodies or chemokine-mediated mobilization of monocytes from the spleen to the heart [12, 24]. However, to our knowledge, B cell-mediated antigen presentation has never been implicated in the pathogenesis of cardiac dysfunction. To further investigate the potential role of MHC II-mediated antigen presentation in ischemic myocardial injury, we isolated MHC II-associated peptides from splenic B cells of sham-operated and ischemic HF mice (4 weeks post-MI) and analyzed them via mass spectrometry (Fig.3e). Peptides enriched more than 1.2-fold in MHC II molecules from post-MI B cells were analyzed *in silico* with a software that predicts peptide binding to the MHC II isoform expressed on C57BL/6 mice. 54 enriched peptides were identified as predicted binders in the *in-silico* analysis, corresponding to 45 proteins. 18% of these proteins were cardiac-specific (Fig.3f). 13 peptides were predicted as strong binders. These included several cardiac-specific proteins such as cardiac α actin (ACTC1), Titin (TTN) and 2 peptides from myosin heavy chain 6 (MYH6) (Fig.3f bottom panel, Supplementary Table 3).

### B cell-specific deletion of MHC II suppresses adverse cardiac remodeling after adoptive transfer of post-MI splenic B cells

To then investigate B cell-dependent MHC II-mediated antigen presentation as a possible mechanistic driver of adverse cardiac remodeling following ischemic myocardial injury, we used transgenic mice with B cell-specific MHC II deletion (*Cd19^tm1(cre)Cgn/-^H2-Ab1^b-^*^tm1Koni/^ *^b-^*^tm1Koni^) [25] and repeated our adoptive transfer experiments (Fig.4a).

**Fig. 4:**
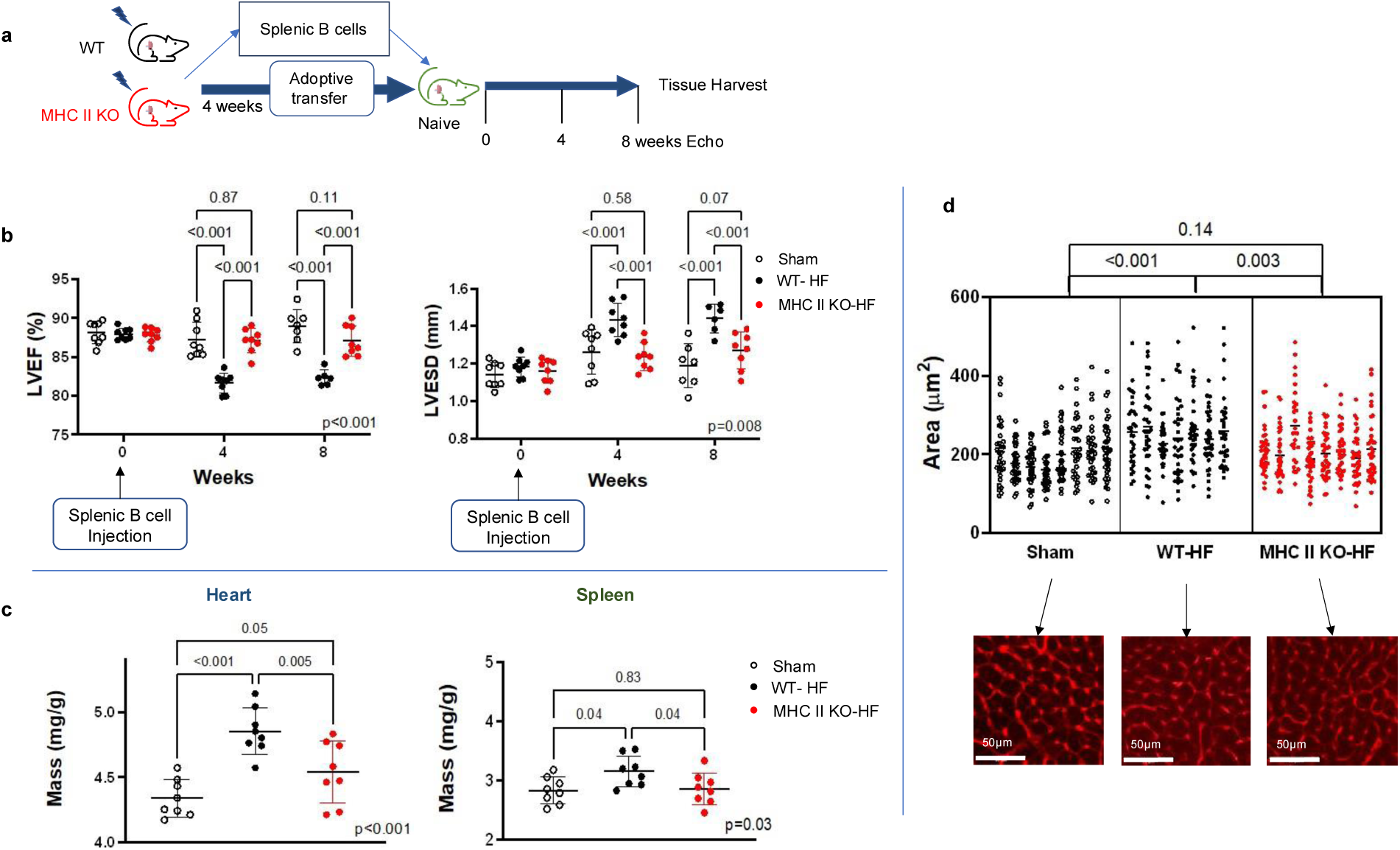
Transfer from ischemic HF mice of splenic B cells deficient in MHC class II does not produce adverse cardiac remodeling in naïve recipient mice at the echocardiographic, gravimetric or histological level. **a**) Schema for adoptive transfer studies of isolated splenic B cells from *Cd19tm1(cre)Cgn/-H2-Ab1b-*tm1Koni/ *b-*tm1Koni mice and wildtype mice following permanent LAD coronary ligation or sham surgery. **b**) Naïve recipients of splenic B cells isolated from HF mice with B cell-specific MHC class II deletion did not experience adverse cardiac remodeling on serial echocardiography over an 8-week period post-adoptive transfer. **c**) Gravimetric data of recipient mice. Recipient mice of B cells deficient in MHCII from HF mice did not experience significant splenic or cardiac remodeling by gravimetric analysis. **d**) Representative images of WGA staining of hearts from recipient mice. Transfer of isolated MHCII-deficient splenic B cells did not induce cardiomyocyte hypertrophy. LAD- left anterior descending. HF-heart failure. n= 8 mice per group. Mean values ± SD are represented.

Four weeks following MI surgery, we performed adoptive transfer of splenic B cells from MHC II-deficient HF mice into naïve wild-type (WT) mice. Flow cytometry analysis confirmed the persistent absence of MHC II on donor B cells at the time of adoptive transfer (Supplementary Fig.11). In a separate experiment, this time using CD45.1 mice as recipients, because the donor MHC II-deficient and WT mice naturally express CD45.2, we confirmed the persistent presence of donor B cells from MHC II-deficient mice 8 weeks following adoptive transfer (Supplementary Fig.12). Adoptive transfer of splenic B cells from WT ischemic HF mice or sham-operated mice served as positive and negative controls, respectively. Splenic B cells isolated from MHC II-deficient mice with ischemic HF did not induce adverse cardiac remodeling upon adoptive transfer as assessed by echocardiography (Fig.4b; mean LVEF 87.10±1.9% vs. 88.96±2.2% MHC II-deficient HF vs. WT sham recipient, q = 0.11 at 8 weeks; mean LVESD 1.27±0.1mm vs. 1.19±0.1mm MHC II-deficient HF vs. WT sham recipient, q = 0.17 at 8 weeks). Gravimetric data revealed borderline increases in cardiac mass and the absence of splenic hypertrophy following the transfer of splenic B cells from MHC II-deficient HF mice (Fig.4c; spleen mean 2.86±0.3 vs. 2.83±0.2 mg/g MHC II-deficient HF vs. WT sham recipient, q = 0.83; heart mean 4.54±0.2 vs. 4.34±0.1 mg/g MHC II-deficient HF vs. WT sham recipient, q = 0.05). Cardiomyocyte hypertrophy was also markedly reduced in recipients of splenic B cells from MHC II-deficient HF mice compared to recipients of splenic B cells from WT HF mice (Fig.4d).

To reinforce the findings from the echocardiographic analysis, we performed proteomics analyses of left ventricular samples from recipients of splenic B cells from WT sham, WT ischemic HF and MHC II-deficient ischemic HF. This analysis confirmed adverse cardiac remodeling in recipients of splenic B cells from WT HF mice relative to recipients from WT sham mice and highlighted blunted adverse cardiac remodeling in recipients of splenic B cells from MHC II-deficient HF mice (Fig.5a-b). There was significant upregulation of key proteins involved in adverse cardiac remodeling (e.g. TPM and TNNT2), cell death and protein turnover (BAG3 and UBB) in recipients of WT HF splenic B cells relative to MHC II-deficient HF splenic B cells. Interestingly, in recipients of MHC II-deficient HF splenic B cells, there was significant upregulation of cardiac repair/stabilization proteins in response to stress like MYOM1, GYG1, CCT2 and NID-1 (Fig.5c).

**Fig. 5:**
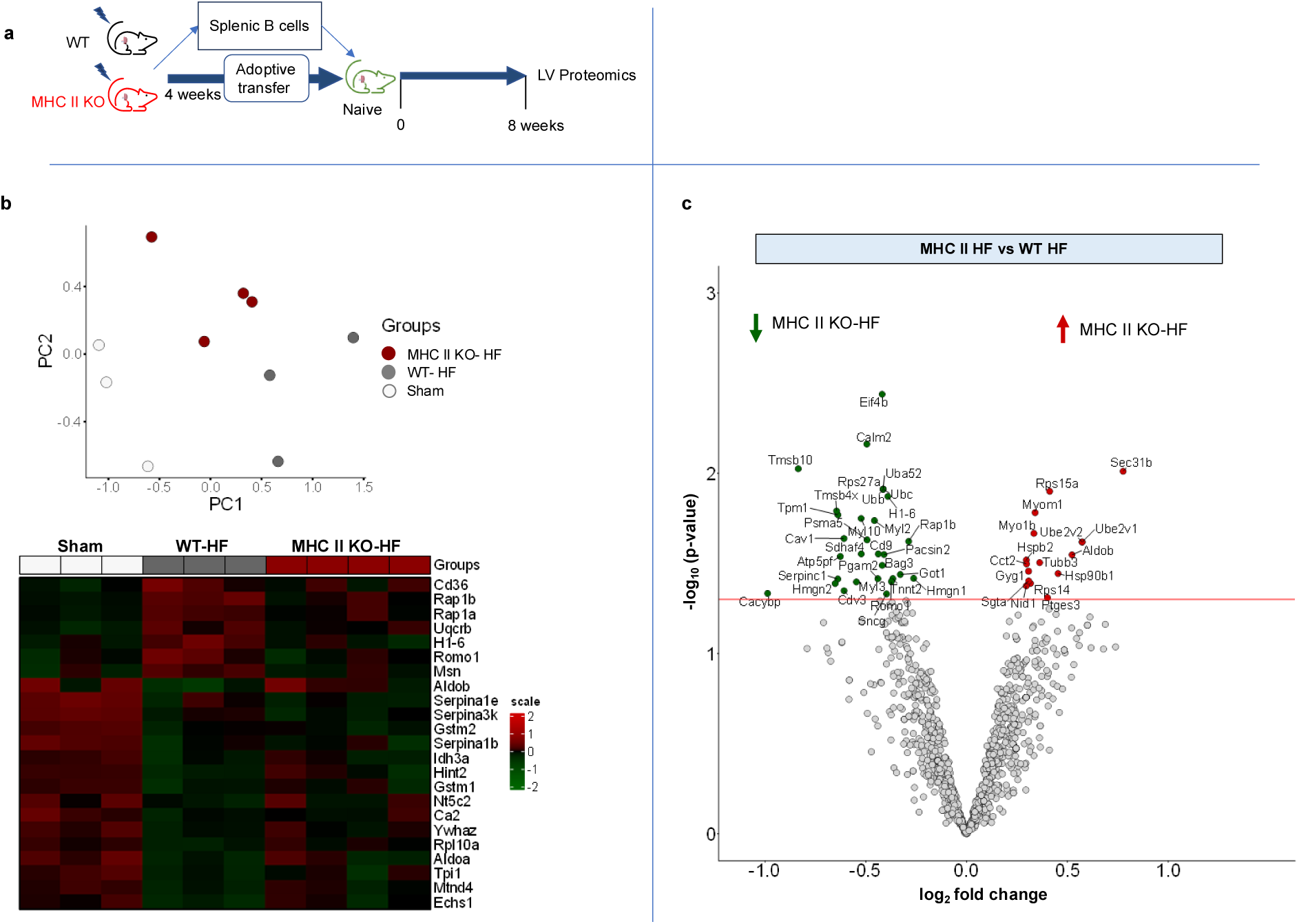
Mass spectrometry of left ventricle shows that transfer of splenic B cells from mice with ischemic HF into naïve recipients results in adverse cardiac remodeling, which is blunted when splenic B cells are deficient in MHC class II. **a**) Schema for adoptive transfer studies of isolated splenic B cells from *Cd19tm1(cre)Cgn/-H2-Ab1b-*tm1Koni/*b-*tm1Koni mice and wildtype (WT) mice following permanent coronary artery ligation or sham surgery. LV-Left ventricle. **b**) PCA plot of protein clusters within each animal across the three groups and heatmap of proteins significantly changed between WT heart failure (WT-HF) and sham. Transfer of isolated MHC II-deficient splenic B cells from heart failure mice (MHC II KO-HF) shifts cardiac proteome of naïve recipients towards control. **c**) Volcano plot of proteomics comparison between WT and MHC II-deficient heart failure recipient mice. Protein markers of adverse cardiac remodeling are down regulated in naïve recipients of heart failure MHC II-deficient splenic B cells relative to wildtype recipients. n=3-4 mice per group. LIMMA statistical test.

### Follicular splenic B cells mediate MHC II-dependent cardiac adverse remodeling post-MI

We ran several control experiments to support proper interpretation of the results obtained using MHC II-deficient B cells. First, we repeated the adoptive transfer experiment utilizing *Cd19^tm1(cre)Cgn/-I^* mice as donors and confirmed that there were no off-target effects of the Cre transgene (i.e. The results from these experiments were comparable to those using WT controls, Supplementary Fig.13).

We then performed immunofluorescence to assess potential changes in serum antibodies targeting myocardial proteins. We incubated heart sections from genetically B-cell deficient μMT mice with serum collected from recipients of isolated splenic B cells from WT sham, WT HF and MHC II-deficient HF mice 8 weeks post adoptive transfer and stained the histological sections with anti IgG and IgM antibodies. We observed no differences in IgG-positive and IgM-positive stain intensities across groups (Supplementary Fig.14).

Finally, we analyzed splenic B cells from MHC II-deficient ischemic HF (4 weeks post- MI) and sham-operated mice. WT HF mice and sham-operated mice served as positive and negative controls, respectively. We found that the deletion of MHC II on B cells was associated with an increase in the percentage of follicular B cells and a decrease in marginal zone B cells post-MI (Supplementary Fig.15). This raised the possibility that the deletion of MHC II on B cells might suppress adverse cardiac remodeling post-adoptive transfer simply due to the decrease in marginal zone B cells. We, therefore, repeated our adoptive transfer experiments, this time transferring either follicular B cells or marginal zone B cells from HF or sham-operated mice (Fig.6a).

**Fig. 6:**
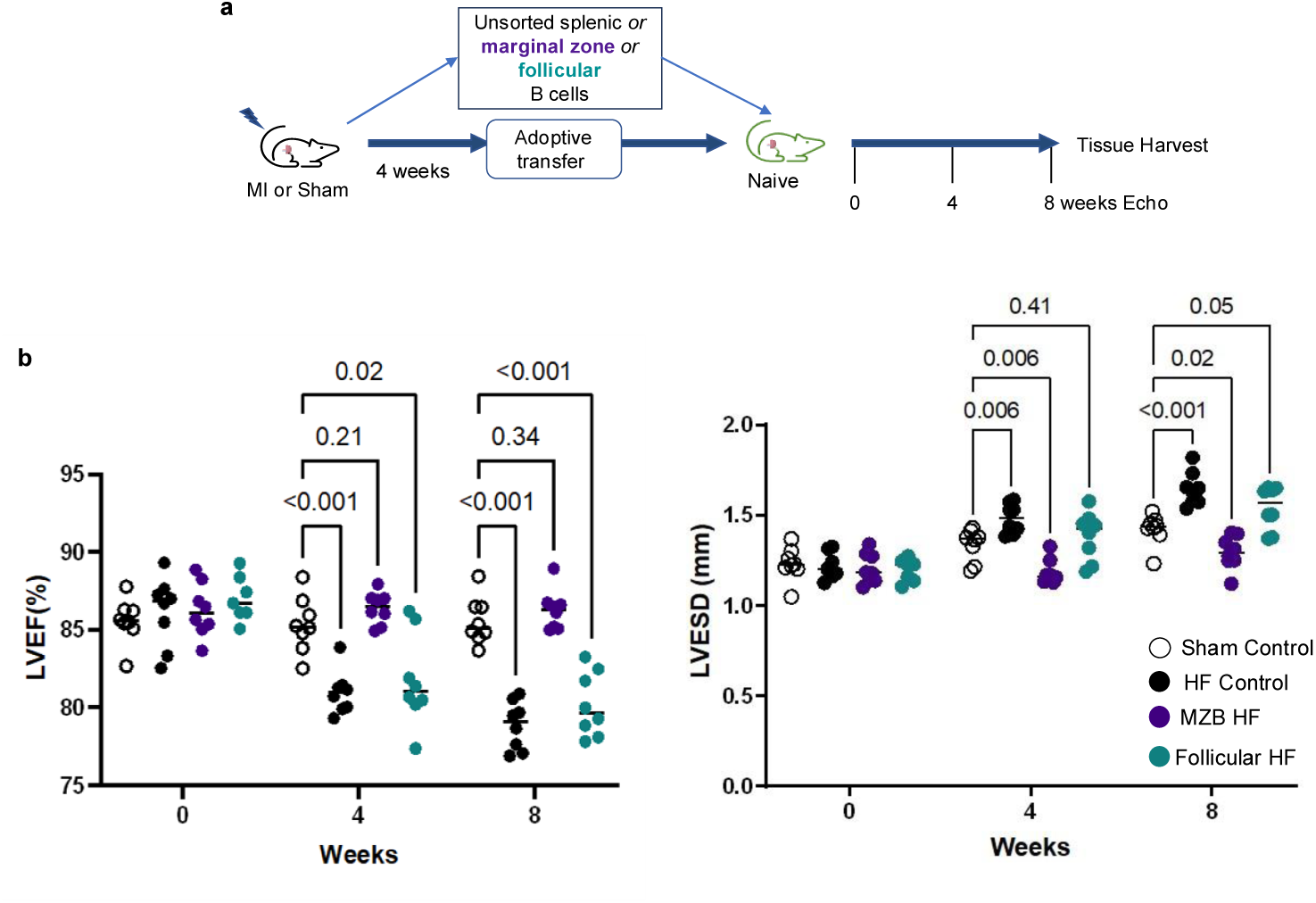
Splenic Follicular B cells promote adverse cardiac remodeling in ischemic HF. **a**) Schema for adoptive transfer studies. Four weeks following permanent LAD coronary ligation, follicular splenic B cells or marginal zone splenic B (MZB) cells were transferred to naïve recipient mice. The adoptive transfer of unsorted splenic B cells from mice 4 weeks following coronary artery ligation (HF control) or sham surgery served as positive and negative controls, respectively. Recipient mice underwent serial transthoracic echocardiography over 8 weeks. **b**) On echocardiography, adoptive transfer of splenic follicular B cells from heart failure (HF) mice resulted in a reduction in left ventricular ejection fraction (LVEF) (p<0.001) and an increase in left ventricular end-systolic diameter (LVESD) in naïve recipient mice over eight weeks (p=0.05) compared to recipients of splenic B cells from sham-operated mice. The adoptive transfer of marginal zone B cells from HF mice did not result in LVEF reduction or LV dilatation. n = 8 mice per group. Mean values ± SD are represented.

Splenic follicular B cells promoted adverse cardiac remodeling in naïve mice, evidenced by reduction in LVEF and increase in LVESD relative to sham (Fig.6b); while marginal zone B cells had no effect (Fig.6b). This identified splenic follicular B cells as the key mediators of adverse cardiac remodeling along the cardio-splenic axis post-MI and reinforced a direct effect of B cell MHC II deletion on B cell-dependent adverse cardiac remodeling post-MI.

### Single cell RNAseq based analysis indicates that adoptively transferred splenic B cells from HF mice modulate myocardial T cells via an MHC-II dependent function

The results presented so far indicate that in ischemic HF, splenic B cells promote adverse cardiac remodeling through an MHC II-dependent function, likely MHC II-mediated antigen presentation. We used scRNA-seq to investigate the effect of adoptive transfer of splenic B cells from HF mice on the cardiac immune cells of recipients. We isolated myocardial immune cells 8 weeks post-adoptive transfer of splenic B cells from WT-HF, MHC II-deficient HF or WT sham operated mice and analyzed them via scRNAseq (Fig.7a-b). We focused our analysis on T cell gene expressions as B cells are known to present antigens to T cells. Comparative gene expression analysis of recipients of splenic B cells from WT-HF vs WT sham operated mice revealed a list of 70 DEGs in myocardial CD4 T cells (12 downregulated, 58 upregulated, adjusted p<0.05) and 745 DEGs in myocardial CD8 T cells (6 downregulated, 739 upregulated, adjusted p<0.05, Supplementary Table 4). When comparing recipients of MHC II-deficient HF splenic B cells to recipients of WT-HF splenic B cells, we found minor differences in CD4 T cells, where 33 DEGs were upregulated and 15 DEGS downregulated (adjusted p<0.05, Fig.7c, left bottom). The difference was more pronounced in CD8 T cells in which only 2 genes were upregulated, and 84 genes were downregulated (Fig.7c, right bottom; Supplementary Table 4). To gain insights into the biological significance of these gene expression changes we used Ingenuity Pathways Analysis (IPA). IPA showed that adoptive transfer of splenic B cells from HF mice resulted in upregulation of interferon gamma (IFNγ) related signaling and other downstream signaling pathways in CD8 T cells (Supplementary Fig.16). Depletion of MHC II from B cells resulted in downregulation of IFNγ and STAT5B related signaling in CD8 T cells (Supplementary Fig.16). In CD4 T cells, IPA showed that deletion of MHC II on B cells resulted in suppression of HSF1 mediated signaling, a signaling pathway that drives the development of regulatory T cells [26].

**Fig. 7:**
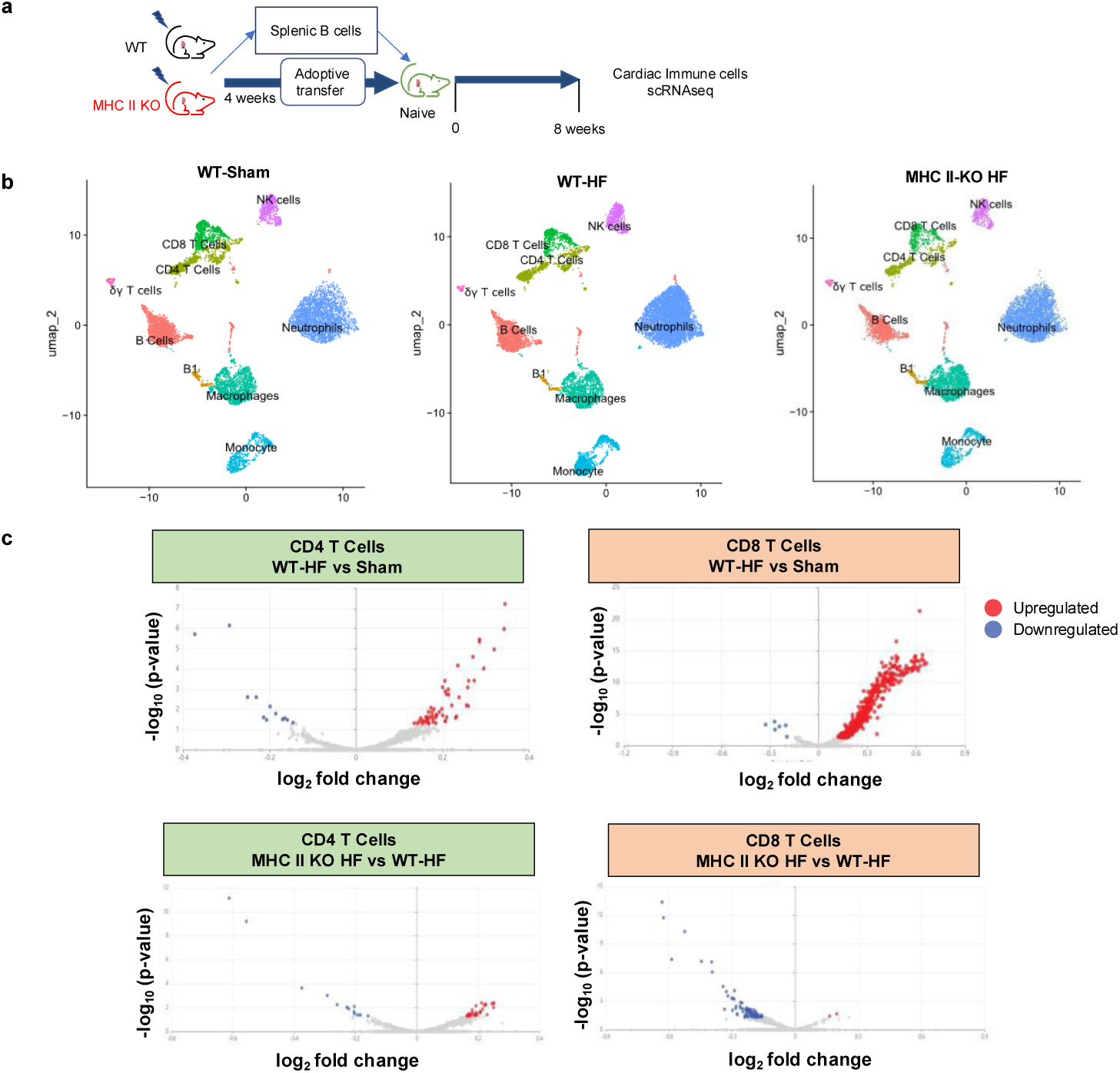
Adoptive transfer of splenic B cells from mice with ischemic HF produces MHC-II dependent changes in myocardial T cells 8 weeks post transfer. **a**) Schema for scRNA-seq based analysis of chronic changes in myocardial immune cells mediated by splenic B cells in the context of ischemic HF. 4 weeks post-MI splenic B cells from WT or *Cd19tm1(cre)Cgn/-H2-Ab1b-*tm1Koni/ *b-*tm1Koni (MHC II KO) mice were isolated and transferred into naïve recipients. 8 weeks later myocardial immune cells were isolated from recipient animals and analyzed via scRNA-seq. **b**) UMAP plots of myocardial immune cells 8 weeks post adoptive transfer of splenic B cells from WT sham, WT HF or MHC II deficient HF mice. **c**) Ingenuity Pathway Analysis (IPA) generated volcano plots of significantly differentially expressed genes; adjusted p<0.05, (n = 6 mice per group). HF= heart failure.

Taken together, these observations suggest that post-MI splenic follicular B cells promote adverse cardiac remodeling in ischemic heart failure through MHC II-dependent modulation of myocardial T cells.

### Ischemic heart failure in humans is associated with differential expression of antigen processing and presentation pathways in B cells

To evaluate the clinical relevance of our observations, we next sought to assess differentially expressed biological pathways in B cells isolated from patients who experienced ischemic myocardial injury. Since our murine data showed that changes in splenic B cells were reflected in peripheral blood B cells (Supplementary Fig.7) and isolation of splenic B cells from patients was not feasible, we performed a focused analysis of publicly available peripheral blood scRNAseq data from patients with ST-elevation myocardial infarction (STEMI) [27] and patients with established ischemic HF relative to healthy patients [28]. In both datasets, we identified peripheral blood B cells and performed differential gene expression analysis.

The STEMI dataset included peripheral blood scRNAseq data of 38 patients 24-hours and 8-weeks post-STEMI in addition to 38 healthy controls [27]. We performed DEG analysis within B cells, comparing post-STEMI patients at both timepoints with healthy controls (Fig.8a). We identified 459 DEGs at 24-hours post-STEMI and 493 DEGs 8-weeks post-STEMI. KEGG pathway analysis of these DEGs highlighted antigen processing and presentation as a differentially regulated pathway both at 24-hours post-STEMI (q = 0.03) and 8-weeks post-STEMI (q = 0.03; Fig.8b), with genes such as HLA-DRB5 and CIITA being differentially expressed and matching this pathway. In the second dataset of three patients with established ischemic cardiomyopathy and one matched healthy control (Fig.8c) [28], we identified 302 DEGs. Antigen processing and presentation was identified here as a top differentially regulated pathway in patients with ischemic HF (q = 2.09E-21; Fig.8d). DEGs relevant to these pathways included HLA-DRB1, HLA-DMA, HLA-DQA1 and -DQA2, HLA-DQB1, which all encode for various MHC II proteins. Complete lists of the DEGs within the antigen processing and presentation pathway are provided in Supplementary Table 5.

**Fig. 8:**
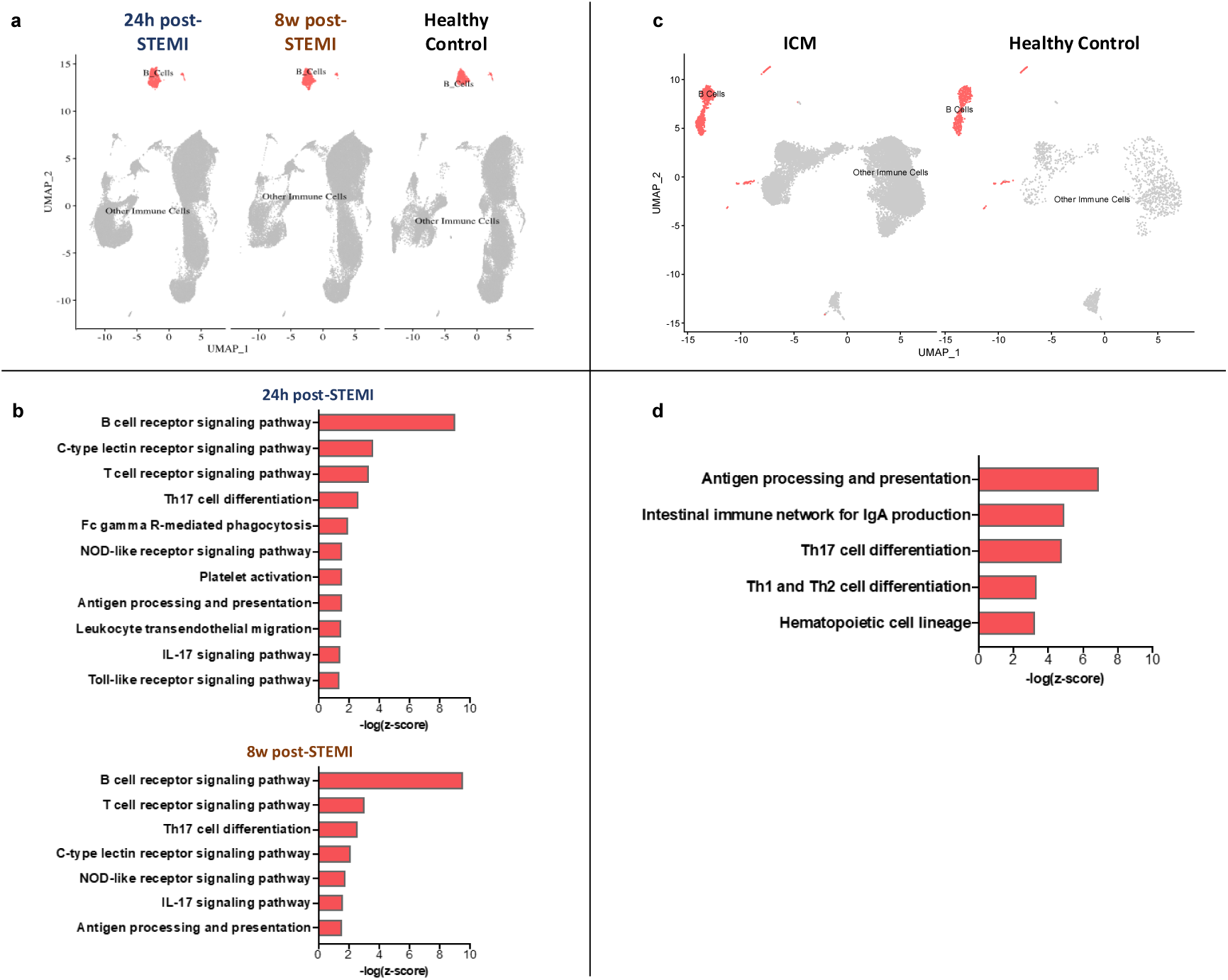
Antigen processing and presentation is differentially expressed in circulating B cells in humans post-STEMI or with ischemic cardiomyopathy. **a**) Single cell sequencing of peripheral blood B cells from 38 patients 8-weeks or 24-hours post-STEMI and 38 healthy controls. UMAP plots are shown with B cells identified. **b**) KEGG pathway analysis of differentially expressed genes (log_2_foldchange≤-1≥1, p<0.05) revealed significant differential regulation of antigen processing and presentation in peripheral B cells of patients at both 24 hours and 8 weeks post STEMI. **c**) Single cell sequencing of peripheral blood B cells from 3 patients with ischemic cardiomyopathy (ICM) and 1 healthy patient. UMAP plots are shown with B cells identified. **d**) KEGG pathway analysis of differentially expressed genes (log_2_foldchange≤-0.58≥0.58, FDR<0.05, p<0.05) revealed significant differential expression of antigen processing and presentation in peripheral blood B cells from ICM patients. All immune system pathways with FDR <0.05 are listed. Th = T helper. NOD = nucleotide oligomerization domain. IL = interleukin.

## Discussion

In this study, we report for the first time that splenic follicular B cells promote adverse cardiac remodeling in ischemic heart failure. We show that splenic B cells present cardiac antigens on MHC II molecules after MI and that their role in promoting adverse cardiac remodeling along the cardio-splenic axis is largely dependent on MHC II. These findings change our understanding of B cell biology in the post-MI period and point to B cell-mediated antigen presentation as a potential novel and specific therapeutic target to prevent and treat ischemic heart failure.

Previous research has established that immune activation along the cardio-splenic axis is critical in ischemic heart failure [7, 8, 29–32]. Although B cells are the most prevalent splenic immune cell across multiple vertebrates, there is limited data on the role of B cells within the cardio-splenic axis. To address this knowledge gap, we employed the adoptive transfer model developed by Ismahil et al., which uniquely allows for the isolation of the functional effects of specific immune cell types within the cardio-splenic axis of ischemic heart failure [8]. The adoptive transfer of splenic B cells from mice with ischemic heart failure resulted in adverse cardiac remodeling with evidence of persistent LV dilatation, reduced LV systolic function, and cardiomyocyte hypertrophy in naïve mice. Our findings thus provide compelling evidence that B cells play a significant role within the cardio-splenic axis of ischemic heart failure.

Our data corroborates previous observations regarding the dynamics of B cell transit to the heart [10], in that adoptively transferred splenic B cells are re-distributed between the heart, spleen, and peripheral blood of recipient mice, suggesting recirculation of transferred splenic B cells between the heart and spleen. This does not exclude distribution to other organs, which has indeed been described in prior studies in naïve mice [10]. The observation that MI triggered the differential regulation of antigen processing and presentation pathways across splenic, myocardial and circulating B cells further supports the notion that B cells actively recirculate between the spleen and the heart through the peripheral blood after MI. Notably, the persistent unique gene expression profile observed in B cells four weeks post-injury suggests that MI induces long-lasting alterations in B cells even in the absence of observable changes in B cell frequency.

Analyzing the MHC-II associated peptidome of post-MI splenic B cells we found enrichment of myocardial antigens. MYH-6, a protein that has been previously described as preferential target of cardiotropic T cells in mice and humans [33, 34] was one of the strong MHC II binders detected, at 2 distinct regions (two different MYH-6 peptides were identified as enriched). We did not observe significant clonal expansion of B cells post-MI. We cannot exclude that this might be the result of insufficient sequencing depth, yet there were no significant changes in the serum concentration of cardiac autoantibodies. Data from the RITA-MI trial also suggests a lack of autoantibody involvement, as serum immunoglobulin levels were not significantly changed from time of MI (baseline) throughout B-cell depletion (Rituximab treatment) [35]. This does therefore present the possibility that splenic B cells may acquire and present cardiac antigens through B cell receptor (BCR)-independent mechanisms rather than through typical BCR engagement [36]. BCR-independent, low-efficiency, antigen uptake has been well documented in B cells [37]. We speculate that after an MI, release of high concentration of cardiac autoantigens might generate appropriate conditions for BCR-independent antigen uptake. Our scRNA data highlights the involvement of pattern recognition receptor (PRR) pathways; thus, there is a possibility that activation of damage-associated molecular patterns (DAMPs) related mechanisms may be the preceding step to loading of cardiac antigens on MHC II molecules of splenic B cells. Further studies will be needed to understand in detail the mechanistic steps that lead to the loading of myocardial antigens on MHC II molecules of splenic B cells.

The findings from our adoptive transfer experiments, which involved post-MI B cells with either intact or deficient MHC II expression, underscore the importance of MHC II in mediating adverse cardiac remodeling along the cardio-splenic axis. Specifically, the absence of MHC II led to a reduction in adverse cardiac remodeling in recipient mice, as assessed by echocardiography, gravimetric analysis, and mass spectrometry of myocardial tissue. While MHC II deletion can influence various B cell functions, including survival, replication, and chemokine production, our results suggest that MHC II-deficient B cells can survive after adoptive transfer and maintain comparable frequencies in recipient animals relative to wild-type B cells. Notably, we found that alterations in the composition of splenic B cells due to MHC II deletion did not account for the phenotype observed in recipients of MHC II-deficient splenic B cells.

We cannot entirely rule out the impact of MHC II deletion on the secretion of B cell-specific chemokines. This is a limitation of our study. Yet, collectively, our findings from the MHC-II associated peptidome analysis and the experiments with MHC-II deficient B cells strongly suggest that splenic B cells modulate adverse cardiac remodeling after MI through MHC II-dependent antigen presentation. It bears emphasis that we noted mild effects of post-MI MHC II-deficient splenic B cells, with trends toward increased cardiac mass, cardiomyocyte cross-sectional area, (Figure 4 c,d) and mild changes at the proteomic level (Figure 5), indicating some degree of adverse cardiac remodeling. This may stem from cytokine-dependent, MHC-II independent, mechanisms by which B cells contribute to cardiac remodeling after MI [12, 38].

B cells typically present antigens via MHC II to CD4 T cells and, under specific conditions, CD8 T cells [39, 40], leading to T cell activation. Prior studies have implicated both CD4 and CD8 T cells in adverse cardiac remodeling following myocardial injury [41–44]. In the present study, we found that the adoptive transfer of splenic B cells from ischemic HF modulated gene expression of both myocardial CD4 T cells and myocardial CD8 T cells in an MHC II dependent manner. In the CD8 compartment, we observed MHC II dependent activation of IFNγ signaling, a key mediator of immune mediated myocardial damage [45–47]. The downregulation of the mediator STAT5B in the absence of MHC II further supports this dependency as altered STAT5B expression and activation has been implicated in various cardiac pathologies [48, 49]. These findings therefore suggest that adoptive transfer of post-MI splenic B cells triggers activation of myocardial T cells, ultimately contributing to adverse cardiac remodeling. Notably, we observed a stronger signal for MHC II dependent CD8 T cells activation, and minor changes in CD4 T cells. It remains unclear whether this finding reflects a cascade of activation initiated by CD4 T cells (with amplification of the activation along the signaling cascade) or if B cells independently engage both CD4 and CD8 T cells, with a stronger effect on CD8 T cells. Further investigations will be needed to clarify this interaction.

Our scRNAseq analyses of peripheral blood B cells from patients with ischemic myocardial injury showed differential regulation of antigen processing and presentation in both the acute and chronic post MI phases. Several genes involved in this pathway showed shared differential expression between mice and humans, including multiple MHC complexes, CD74 (which stabilizes MHC II molecules) [50], and HSPs, which are critical modulators of effector response post antigen presentation [51, 52].This is noteworthy, as it points to a translational relevance of our findings. However further work will be needed to assess the direct involvement of these conserved genes and the extent to which B cell-mediated antigen presentation modulates adverse cardiac remodeling in patients who experience an MI.

In summary, our findings reveal that MI triggers long-lasting changes in splenic B cells; that splenic B cells present myocardial antigens via MHC II following myocardial infarction; and that B cells circulating between the spleen and the heart drive post-MI adverse cardiac remodeling through MHC II-mediated mechanisms. These observations not only expand our current understanding of B cell biology but also reshape existing models of immune-cardiac interactions post-myocardial injury [2, 53–55]. Moreover, they highlight the potential of targeting MHC II-dependent B cell functions as a novel therapeutic approach to prevent and treat ischemic heart failure. Further studies will be needed to understand whether our findings are relevant to other conditions characterized by acute or chronic cardiac damage.

## Methods

All studies were approved by the Institutional Animal Care and Use Committee (ACUC) of the Johns Hopkins University School of Medicine, under protocol numbers MO20M388 and M023M238.

### Mouse Models and Surgical Protocol

Male 10-14- week-old mice were used for all experiments unless otherwise specified. All mice were purchased from Jackson Laboratory (Bar Harbor, ME) or bred in the Johns Hopkins vivarium facility. The following strains were used: C57BL/6J strain N. 000664, B6.SJL-*Ptprc^a^ Pepc^b^*/BoyJ (CD45.1) strain N. 002014, B6.129P2(C)-*Cd19^tm1(cre)Cgn^*/J strain N. 006785, and B6.129X1-*H2-Ab1^b-tm1Koni^*/J strain N. 013181. To generate mice with B cell-specific MHC class II deletion, B6.129P2(C)-*Cd19^tm1(cre)Cgn^*/J and B6.129X1-*H2-Ab1^b-tm1Koni^*/J mice were bred for target *Cd19^tm1(cre)Cgn/-^H2-Ab1^b-^*^tm1Koni/*b-*tm1Koni^ mice. Mice were randomly grouped into sham donors or HF donors. HF donors underwent left thoracotomy with permanent ligation of the left anterior descending (LAD) coronary artery to induce an MI. For the desired condition of ischemic HF, mice were utilized at four weeks post- MI surgery [56, 57]. Sham donors underwent sham intervention. Experiments referring to wildtype mice include a mix of heterozygous *Cd19^tm1(cre)Cgn/-^H2-Ab^-^*^/*-*^, homozygous *Cd19^tm1(cre)Cgn/-^* littermate controls and syngeneic C57BL/6J strain N. 000664.

### Adoptive Transfer Studies

Mice were euthanized four weeks following MI or sham surgery. Animals were gently perfused through heart apex with 3mL Hanks’ Balanced salt solution (HBSS) without CaCl_2_, MgCl_2_ or MgSO_4_ (GIBCO). Spleens were harvested and placed into a 35mm tissue culture dish with phosphate-buffered saline (PBS). Splenocytes were isolated as previously published by Ismahil et al [8]. Briefly, the spleen was minced finely using scissors and the end of a syringe plunger. The solution was then pipetted through a 40μm cell strainer into a sterile 50mL tube. The tube was centrifuged at 250g for 5 minutes at 4°C. After decanting the supernatant, the pellet was resuspended with 5mL of ACK lysis buffer (Quality Biological) for 5 minutes at room temperature to remove red blood cells (RBCs). After 5 minutes of lysis, 30mL of PBS was added to each tube and each tube was centrifuged at 250g for 5 minutes at 4°C. The supernatant was decanted, and pelleted splenocytes were then resuspended in sterile 0.9% sodium chloride for a final concentration of ∼1×10^8^ total splenocytes in 150μL per retro-orbital injection into naïve recipient mice.

For adoptive transfer studies with purified B cells, splenic B cells were isolated according to the MojoSort™ Mouse Pan B Cell Isolation Kit protocol (BioLegend), with slight modifications. Briefly, after mincing and filtering each spleen as described above, the pellet of each spleen was resuspended with 4mL of Mojo buffer and then filtered again into fluorescence-activated cell sorting (FACS) tubes. Each tube containing cells from one spleen was then centrifuged at 250g for 5 minutes at 4°C. After decanting the supernatant, the pellet was resuspended with cold 2mL Mojo buffer. Samples were then incubated in 200μL of primary antibody cocktail for 15 minutes on ice, followed by another 15 minutes incubation with 200μL of Nanobeads. Incubation was stopped with 1.5mL Mojo buffer, and magnet sorted for 5 minutes. The cell suspension was decanted into a fresh FACS tube, magnet incubation was repeated to increase the yield of isolated B cells. The suspension of each spleen was then centrifuged, and each pellet was resuspended in FACS buffer. The splenic B cell solutions from mice of the same experimental group were combined into a single tube and then centrifuged prior to final resuspension in sterile 0.9% sodium chloride for retro-orbital injection of ∼1×10^7^ splenic B cells in 150µL into naïve recipient mice. A small sample was obtained pre- and post-antibody incubation to evaluate purity and yield by flow cytometry (Supplementary Fig.1). Purified splenic B cells consistently had >95% purity.

For adoptive transfer studies with splenic B cell subtypes, single cell suspensions were prepared as above and then stained with CD19-BV421, CD21-APC, and CD23-PE antibodies and propidium iodide. Using MoFlo sorter, marginal zone B (MZB) cells and follicular B cells were sorted as CD19^+^/CD21^+^/CD23^low^ and CD19^+^/CD21^low^/CD23^high^, respectively. Following collection, samples from different mice were pooled, and all sorted and approximately 1.5×10^6^ MZB cells and 1×10^7^ follicular B cells were injected into naïve recipient mice, maintaining the proportion of these two cell types in unfractionated splenocytes. These numbers of splenic B cell subtypes were representative of the proportion relative to the total number of unsorted B cells from a mouse spleen.

### Echocardiography

Mouse transthoracic echocardiography was performed in conscious mice prior to retro-orbital injection to assess baseline cardiac structure and function, and again 4-, 8- and 16- weeks post adoptive transfer. Echocardiography was performed using a VisualSonics Vevo 2100 Imaging System. M-mode parasternal short images were obtained. LVEF and LVESD were measured using Vevo Lab software v5.8.2.

### Gravimetric and Histological Analyses

Eight weeks after the adoptive transfer, recipient mice were euthanized, and hearts and spleens were dissected and weighed. Tibia length and body weight were recorded to normalize the weights of each mouse heart and spleen. Heart tissue was fixed with 10% neutral buffered formalin, - embedded in paraffin, and stained with WGA using standard protocols.

### Isolation of Myocardial Mononuclear Cells

To analyze myocardial immune cells via flow cytometry, the heart was digested as previously described with slight modifications [58]. Each heart was perfused with 3mL of Hank’s Buffered Salt Solution (HBSS) with calcium and magnesium infused into the left ventricle over 1 minute with aortic clamping. Hearts were then dissected and cut transversely, approximately at the mid-papillary muscle level, or approximately ∼5mm below the atrioventricular groove. The superior section was used for flow cytometry, and the apical section was used for histological analysis. The heart tissue for flow cytometry was finely minced and 60mg was suspended in 3mL HBSS. Heart tissue was digested with 300U DNAse, 625U collagenase II, and 50U hyaluronidase shaken for 30 minutes at 300rpm and 37°C. At the end of the 30 minutes, additional HBSS was added to each tube to neutralize the enzymatic digestion. Each tube was then centrifuged at 250g for 5 min at 4°C. The supernatant was decanted, and the pellet was resuspended in 5mL of ACK lysis buffer for 5 minutes at room temperature. The suspension was then diluted with PBS and filtered through a 40μm strainer. Each cell suspension was centrifuged at 250g for 5 min prior to resuspension in FACS buffer and filtering through a 35μm mesh strainer into a sterile FACS tube.

### Isolation of Splenocytes for Flow Cytometry

The spleens were removed and prepared as described above for adoptive transfer of splenocytes. Following completion of RBC lysis, cells were resuspended in FACS buffer and filtered through 35μm cell strainers into sterile FACS tubes.

### Isolation of Peripheral Blood Cells for Flow Cytometry

Peripheral blood (∼100μL) was collected via venipuncture of the facial vein into 1.5mL Eppendorf tubes containing 100μL heparin. RBCs were lysed with 1mL ACK buffer for 5 minutes at room temperature and then centrifuged at 250g for 5 minutes at 4°C. Cells were resuspended in 500μL FACS buffer then filtered through 35μm cell strainers into FACS tubes.

### Flow Cytometry

Isolated cell suspensions were stained for 30 minutes on ice with the following fluorescently conjugated antibodies: CD3-APC/Fire810 (clone 17A2, BioLegend), PD-1-BUV615 (clone 29F.1A12, BD Horizon), CD44-BV737 (clone IM7, Invitrogen), CXCR3-BV421 (clone CXCR3-173, BioLegend), CD62L-BV510 (clone MEL-14, BioLegend), CD8 (clone 53-6.7, BioLegend), CD69-BV650 (clone H1.2F3, BioLegend), CD27-BV785 (clone LG.3A10, BioLegend), KLRG-1-PerCP/Cy5.5 (clone 2F1/KLRG1, BioLegend), CD103-PerCP/eFluor710 (clone 2E7, Invitrogen), CXCR6-PE (cloneSA051D1, BioLegend), CD127-PE/Dazzle594 (clone A7R34, BioLegend), ICOS-PE/Cy5 (clone 15F9, BioLegend), Ki-67-AF488 (clone SolA15, Invitrogen), Granzyme B-Pacific Blue (clone GB11, BioLegend) CD25-BUV395 (clone PC61, BioLegend), CD4-BV650 (clone GK1.5, BioLegend), CCR6-BV785 (clone29-2L7, BioLegend), CXCR5-PerCP/eFluor710 (clone SPRCL5, Invitrogen), FOXP3-AF488 (clone MF-14, BioLegend), CTLA-4-PE (clone UC10-4B9, BioLegend), T-bet-PE/Dazzle594 (clone 4B10, BioLegend), RORrt-PE/Cy7 (clone Q31-378, BD Pharmingen) CD45-BUV395 (clone 30-F11, BD Horizon), CD73-BV421 (clone TY/11.8, BioLegend), GL7-Pacific Blue (clone GL7, BioLegend), IgM-BV605 (clone RMM-1, BioLegend), CD80-BV650 (clone 16-10A1, BioLegend), CD21/CD35-BV711 (clone 7E9, BioLegend), CD19-BV750 (clone 6D5, BioLegend), I-ab-AF488 (clone KH-74, BioLegend), CD5-PerCP (clone 53-7.3, BioLegend), CD1d-PerCP/Cy5.5 (clone L363, BioLegend), CD23-PE (clone B3B4, BioLegend), CD86-PE/Dazzle594 (clone GL-1, BioLegend), CD38-PE/Fire640 (clone 90, BioLegend), CD11b-PE/Cy5 (clone M1/70, BioLegend), PD-L2-PE/Cy7 (clone TY25, BioLegend), CD138-PE/Fire810 (clone 281-2, BioLegend), CD93-APC (clone AA4.1, BioLegend), IgD-APC/Fire 750 (clone 11-26c.2a, BioLegend) CD3-FITC (clone 17A2, Invitrogen), CD3-BV605 (17A2, BioLegend), CD19-BV421 (6D5, BioLegend), CD4-Spark NIR 685 (GK1.5, BioLegend), CD8-BV785 (53-6.7, BioLegend), CD45-PerCP/Cyanine5.5 (30-F11, BioLegend), CCR2-APC (QA18A56, BioLegend), CD64-PE/Cy7 (X54-5/7.1, BioLegend), Ly6C-BV650 (HK1.4, BioLegend), MHCII-BV711 (M5/114.15.2, BioLegend), CD11b-Alexa Fluor 700 (M1/70, BioLegend), Ly6G-APC/Cy7 (1A8, BioLegend), CD45.1-PE (A20, BioLegend). All antibodies were added at 1:1000 dilution for heart and spleen flow cytometry and 1:500 for peripheral blood flow cytometry, with the following exceptions. PD-1 antibody was added at 1:1600 dilution. Ki-67, CXCR6, CD21/CD35, CD19, I-ab, and CD38 antibodies were added at 1:800 dilution. CD44, CD27, CD25, CTLA-4, and T-bet, PD-L2 antibodies were added at 1:100 dilution. Dead cells were excluded using the Zombie NIR dye (BioLegend) at 1:5000 dilution. Flow cytometry was performed on Cytek Aurora Full Spectrum Flow Cytometric Analyzer. Compensation controls were generated using single-color control samples and UltraComp eBeads (Invitrogen). Representative gating strategies are shown in individual figures.

### Single Cell RNA (scRNA) Sequencing

C57BL/6J mice 14-18 weeks-old underwent left thoracotomy with either sham surgery (n = 5) or permanent coronary LAD ligation (n = 11). Blood, heart, and spleen were collected 4 weeks post-sham or MI surgery (“heart failure”; n = 6) or 4 days post-MI surgery (“recent MI”; n = 5). For myocardial immune cell scRNA seq, recipient hearts (n=6/group) were collected 8 weeks after retro-orbital injections of splenic B cells from sham, HF or MHC II KO HF mice; half the heart, cut along the longitudinal axis were further processed for scRNAseq, remnant stored at -80°C. Single-cell suspensions for blood, heart, and spleen were prepared as described above. Heart and spleen samples were stained with 2 drops of Vibrant Dye Cycle Violet Ready Flow Reagent (DCV-Invitrogen) for 30 minutes, shaking at 300rpm and 37°C. Samples were stained with propidium iodide (PI), CD45-PE antibodies for all experiments and CD19-APC antibody for specific B cell subtypes, for 30 minutes on ice. Hashtag antibodies mapped to each experimental group were also added to cell suspensions at a concentration of 2μL per tube. B cells or CD45^+^ cells were sorted using a MoFlo sorter. Equal numbers of the three experimental groups were mixed to reach a total of 20,000-30,000 cells that were combined in equal proportions within the same single cell library. Three libraries were prepared for the post-MI surgery experiment: 1) peripheral blood B cells; 2) splenic B cells; and 3) myocardial B cells. One library was prepared for the myocardial immune cell experiment.

Demultiplexing, barcode processing, single cell 5’ gene counting, and V(D)J transcript sequence assembly and annotation were performed using Cell Ranger (10x Genomics). Gene specific analysis was performed using *Partek™ Flow™* software (v.11.0) for the post-MI surgery experiment and R (v.4.0.2) for myocardial immune cells experiment.

Uniform manifold approximation and projection (UMAP) plots were created on R (v.4.0.2) using Seurat (v.4.1.1) [59] standard parameters unless otherwise noted. Cells were filtered based on their distributions to only include those with 200-3200 unique features for blood, 200-3800 for heart (post-MI experiment), 200-4800 for heart (myocardial immune experiment), and 200-3500 for spleen, and < 8% mitochondrial read counts. Doublets not identified by hashtag demultiplex were removed using scDblFinder (v. 1.24.0). Counts were then logNormalized and scaled, and the top 2000 (post-MI experiment) and top 3000 (myocardial immune experiment) most variable genes were identified using the FindVariableFeatures function. Principal component (PC) analysis was performed using RunPCA and the first 15 PCs (post-MI) and 30 PCs (myocardial immune) were used to find the shared nearest neighbors (SNN) using the FindNeighbours function. Clustering was performed using the FindClusters function with a resolution of 0.2 and visualized using a UMAP projection. B cell and myocardial immune cell subtypes were classified using ScType [18] by using a modified list of known murine B cell and myocardial immune markers (Supplementary Table 6) and mapping mouse gene symbols to their human counterpart using biomart to be compatible with ScType. Analyses of differential expressed genes between conditions per each cell type were performed using DEseq2 (1.46.0), with false-discovery rate (FDR) step-up used for multiple test corrections. Genes with low expression (<10 cells with 1 count) were filtered out prior to model fitting. For post-MI experiments lists of genes that exhibited differential expression based on the threshold of raw p-values<0.05 and log_2_fold change >1 for upregulation of <-1 for downregulation were compiled and then analyzed against the Kyoto Encyclopedia of Genes and Genomes (KEGG) database on the Gene Set Enrichment Analysis (GSEA) website (https://www.gsea-msigdb.org; [19, 20]). Pathways were filtered using KEGG pathway annotations for immune-system specificity and were considered as differentially regulated if q-value from FDR was ≥ 0.05. For the myocardial immune experiment DEGs were filtered using threshold of adjusted p<0.05 and log_2_fold change of ≥0.02 for upregulation and ≤-0.02 for downregulation. The functional analyses and networks were generated using QIAGEN IPA and IPA Interpret (QIAGEN Inc., https://digitalinsights.qiagen.com/IPA and https://digitalinsights.qiagen.com/products-overview/discovery-insights-portfolio/analysis-and-visualization/qiagen-ipa/qiagen-ipa-interpret/) [60].

The Chromium Next GEM Single Cell 5’ Kit v2 (10x Genomics) was used to perform single cell transcriptional analyses. The Chromium Next GEM Single Cell V(D)J Kit was used to assess B cell receptor (BCR) cDNA and clonality. The Chromium Next GEM Single Cell V(D)J Kit was used to obtain BCR cDNA. Full-length and productive contigs were analyzed using scRepertoire (v.2.2.1). Only B cells with fully annotated heavy and light chains were kept for clonotype analysis. CDR3 nucleotide sequences plus V gene were used to define clonotypes. BCRs with >85% normalized Levenshtein distance of the nucleotide sequence were considered clones. All codes used for analyses is publicly available: https://github.com/caro471/Adamo_collaboration_2026.

### Human scRNA Sequencing Analysis

Publicly available scRNA sequencing data of peripheral blood from two cohorts were analyzed: three patients with ischemic cardiomyopathy (ICM) EGA accession: <u>EGAS00001006330</u> [28] and 38 patients 24 hours and 8 weeks following ST segment elevation MI (STEMI) EGA accession: EGAD00001010064 [27] relative to healthy controls. These data were processed on R (v.4.0.2) using Seurat (v.4.1.1) as described above with the following modifications: Cells were filtered based on their distributions to only include those with 200-8000 unique features and <5% mitochondrial reads for the ICM dataset, 200 – 4500 unique features and <12% mitochondrial reads for the STEMI v2 dataset, 200 – 3500 unique features and <8% mitochondrial reads for the STEMI v3 dataset. 3000 variable features were used for both human datasets. Both datasets had notable batch effects that were corrected using Harmony [61]. For both datasets, the first 30 PCs were used for the SNN and clustering. A clustering resolution of 0.5 and 0.4 was used for the ICM and STEMI datasets respectively. B cells were identified using known human B cell-specific makers as previously described (Supplementary Table 6) [62]. For the larger dataset, B cells were pseudobulked per patient and differential gene expression was performed using DEseq2 controlling for sex and 10x chemistry in the regression model. Subsequent KEGG pathway analyses were performed as described above. For the smaller dataset, the defined threshold used was p-value <0.05, FDR step up <0.05and log_2_fold change threshold of ≤-0.58 or ≥0.58 with p<0.05. For the larger dataset, a more stringent log_2_fold change threshold of ≤-1 or ≥1 was used with p-value<0.05.

### Peptidome Analysis

Wildtype (WT) C57BL/6J and *Cd19^tm1(cre)Cgn/-^H2-Ab1^b-^*^tm1Koni/^ *^b-^*^tm1Koni^ (MHC II-KO) mice (n = 16 total) underwent left thoracotomy with either sham surgery (n = 8) or permanent coronary ligation (n = 8). MHC II KO mice were included to serve as negative controls. Four weeks following surgery, spleens were collected from mice and splenic B cells were isolated, snap frozen in liquid nitrogen, and stored at -80°C. Four spleens from the same experimental condition were combined to get 2×10^6^ million B cells per experimental condition. Frozen splenic B cells were lysed and purified for MHC-II bound peptides as previously described [63]. Briefly cells were lysed in 50mM Tris- 150mM NaCl buffer, supplemented with protease inhibitor cocktail (Roche) and 0.5% NP-40 using a bead homogenizer. Lysates were cleared by centrifugation at 2000g and 4°C for 10 minutes, the supernatant was collected and cleared again by ultra-centrifugation at 100,000g and 4°C for 75 minutes. The supernatant was further cleared by flowing through Protein A/G Sepharose column and then incubated for 2 hours at 4°C in Protein A/G Sepharose column cross-linked with MHC-II mAb (M5/114.15.2, BioLegend) with slow agitation (60-80 rpm). Afterwards flowthrough was decanted, and the peptide-bound column was washed four times: first with a buffer containing, 0.05% NP-40, 50mM Tris, 150mM NaCl, 5mM EDTA, 0.1mM PMSF and 1mM Pepstatin A; second with 50mM Tris and 150mM NaCl; third 50mM Tris and 450mM NaCl and finally 50mM Tris. Immuno-bound MHC molecules were then eluted with 10% acetic acid, lyophilized and shipped to Dr Sidoli’s lab and analyzed on an Orbitrap Fusion Lumos (Thermo Scientific) mass spectrometer (details below).

Peptides abundances from WT groups were corrected using peptide abundances from MHC II-KO mice, to eliminate background noise. The average abundance of corrected values was calculated across technical replicates for each peptide. Peptides with mean corrected values above zero were included in further analyses. The fold enrichment for each peptide was determined by dividing values of HF group by sham. The binding affinity (BA) of these peptides were then determined using the NetMHCIIpan-4.3 machine learning tool [64, 65]. Peptides with a fold enrichment of greater than 1.2 and amino acid (AA) sequence length of 9 AAs were submitted into the BA server. Briefly the server is trained on an extensive dataset of naturally occurring peptides covering 650,000 measurements of Binding Affinity and Eluted Ligand Mass spectrometry. The server ranks each peptide according to predicted binding affinity in nanomolar IC50 compared against a set of 100,000 random natural peptides [64, 65].

### Cardiac proteomic sample preparation and analyses

Snap frozen left ventricular tissue collected from recipients of adoptive transfer studies were processed for mass spectrometry analyses using PreOmics® iST-NHS kit according to manufacturer’s instructions. Briefly, tissue was homogenized in LYSE-NHS buffer, sonicated and boiled at 95°C for 10 minutes. Homogenates were cleared by centrifugation at 14,000g for 14 minutes, supernatant was collected, and samples were incubated in DIGEST buffer at 37°C for 1-2 hours. Following digest, samples were labelled using TMTpro^TM^ 16plex Label kit (Thermo Fisher), washed, eluted, lyophilized and processed on an Orbitrap Fusion Lumos (Thermo Scientific) mass spectrometer (details below).

Spectral TMT reporter ion intensities were quantified using the median sweep normalization strategy originally described by Herbrich et al [66], following the implementation reported by Foster et al [67], with minor modifications. Briefly, TMT reporter ion intensities were (1) log₂-transformed, (2) median-centered within each channel, followed by (3) median-centering of each individual spectrum across channels, and (4) protein abundance was determined by median summarization, defined as the median of the log₂-transformed, median-centered intensities from all spectra assigned to that protein within a given channel. Differential protein abundance between experimental groups was evaluated using a Bayesian statistical framework implemented in linear models for microarray data (LIMMA) with multigroup comparisons [68]. Pairwise contrasts were also performed. Moderated p-values were adjusted for multiple testing using the positive false discovery rate (q-value) approach described by Storey & Tibshirani [69]. Heat maps were generated using the ComplexHeatmap R package with hierarchical clustering (Euclidean distance, complete linkage). Principal component analysis (PCA) was performed using the PCAtools R package. All codes used for statistical analysis, including PCA and heat map generation, is publicly available at: https://github.com/Frostman300/MHCII_upload.

### Mass Spectrometry

#### Sample Desalting

Prior to mass spectrometry analysis of both TMT-labeled samples and peptides, samples were desalted using a 96-well plate filter (Orochem) packed with 1 mg of Oasis HLB C-18 resin (Waters). Briefly, the samples were resuspended in 100 µl of 0.1% TFA and loaded onto the HLB resin, which was previously equilibrated using 100 µl of the same buffer. After washing with 100 µl of 0.1% TFA, the samples were eluted with a buffer containing 70 µl of 60% acetonitrile and 0.1% TFA and then dried in a vacuum centrifuge.

#### LC-MS/MS Acquisition and Analysis

Samples were resuspended in 10 µl of 0.1% TFA and loaded onto a Dionex RSLC Ultimate 300 (Thermo Scientific), coupled online with an Orbitrap Fusion Lumos (Thermo Scientific). Chromatographic separation was performed with a two-column system, consisting of a C-18 trap cartridge (300 µm ID, 5 mm length) and a picofrit analytical column (75 µm ID, 25 cm length) packed in-house with reversed-phase Repro-Sil Pur C18-AQ 3 µm resin. Peptides were separated using a 90 min gradient from 4-30% buffer B (buffer A: 0.1% formic acid, buffer B: 80% acetonitrile + 0.1% formic acid) at a flow rate of 300 nl/min. The mass spectrometer was set to acquire spectra in a data-dependent acquisition (DDA) mode from TMT-labeled samples and data-independent mode (DIA) for peptidome samples. Briefly, the full MS scan was set to 300-1200 m/z in the orbitrap with a resolution of 120,000 (at 200 m/z) and an AGC target of 5×10e5. MS/MS was performed in the Orbitrap using the top speed mode (2 secs), an AGC target of 2×10e5 and an HCD collision energy of 45. Search tolerance for MS1 was 10ppm and 20ppm for MS2, fixed modifications N-terminus TMT, lysine TMT and cysteine carbamidomethyl and methionine oxidation set for variable modifications.

Proteome raw files were searched using Proteome Discoverer software (v2.5, Thermo Scientific) using SEQUEST search engine and the SwissProt mouse database (updated April 2023). Any peptide cleavage was specified for digestion, so that the software considered all protein fragments. FDR determined by percolator validator. The mass spectrometry proteomics data have been deposited to the ProteomeXchange Consortium via the PRIDE [70] partner repository with the dataset identifier PXD073894

#### Immunofluorescence staining

Endpoint serum from recipients of adoptive transfer studies were evaluated for cardiac autoantibodies as previously described [71]. Briefly paraffin embedded cardiac cross-sections from congenital B cell-deficient µMT mice were rehydrated and prepared using Tris-EDTA antigen retrieval and incubated with recipient-serum (1:20 dilution) overnight at 4°C. Sections were then washed and autoantibodies against cardiac antigens were then probed for using Biotin-goat anti mouse IgG (clone Poly4053, BioLegend) and rat anti-mouse IgM (clone RMM-1, BioLegend) at 1:1000 dilution. Following washes, the antibodies were detected using streptavidin-Alexa555 (Invitrogen) and anti-rat-Alexa647 (Life Technologies); cardiomyocyte basement membrane was stained with WGA-Alexa488 and nuclei stained with DAPI. Fluorescent images were acquired using the PhenoImager™ HT 3.0, Vectra Polaris (Akoya Biosciences) slide scanner at 40x objective, with the same exposure time. Images were processed using QuPath and analyzed with ImageJ, three ROIs were taken for each section and two sections per sample. Aged mouse serum was used for positive control and negative control were section without serum.

#### Statistical Analysis

Data was initially processed on Microsoft Excel (Microsoft 365). After assessing normality with the Shapiro-Wilk test, Mann-Whitney or unpaired two-tailed t-tests were used for non-parametric and parametric comparisons, respectively, of two groups. To compare data across three or more groups, one-way analysis of variance (ANOVA) or Kruskal- Wallis tests were used for normally distributed or non-Gaussian distributed data, respectively, followed by pairwise comparisons. Serial echocardiographic data were analyzed using two-way ANOVA or mixed-effects analysis followed by pairwise comparisons. WGA data were analyzed using nested t-test for two groups or nested one-way ANOVA followed by pairwise multiple comparisons for three groups. All pairwise multiple comparison tests were corrected using the original FDR method of Benjamini and Hochberg. Outliers were determined using the ROUT method (Q=1%). All statistical tests were performed on GraphPad Prism version 10.6.1 for Windows, GraphPad Software, Boston, Massachusetts USA, www.graphpad.com.

## Acknowledgments

The authors acknowledge the support for the Ross Flow Cytometry core funded by NIH Grant S10OD026859. The authors would also like to thank Dr. Edward Pearce for critical discussions and feedback on the manuscript and Dr. Hao Zhang from the Bloomberg FACS sorting Facility for technical support with FACS sorting. The Sidoli lab gratefully acknowledges for funding the Hevolution Foundation (AFAR), the Einstein-Rockefeller-CUNY-Mount Sinai Center for AIDS Research, the NIH Office of the Director (S10OD030286), and the Aging Biology Foundation.

## Author contributions

OVE and JPL designed experiments, performed experiments, analyzed data, and wrote the manuscript. CD assisted with the analysis of murine and human scRNA data. SR, AB, YS, SS and KK assisted with collection and analysis of experimental data. KB, CC, SS, DBF, MM and SK helped with experimental design, review of data and manuscript edits. LA conceived the project, supervised experimental design and analyses, and wrote the manuscript with OVE and JPL.

## Notes

### Competing Interest Statement

The authors have declared no competing interest.

### Summary of Updates

New data presented in Figures 3, 5, 6 and 7; Results section updated to include new data presented; Discussion section updated; Author list updated.

