## Extended Data for "Splenic follicular B cells promote adverse cardiac remodeling after myocardial infarction via MHC II-dependent antigen presentation"

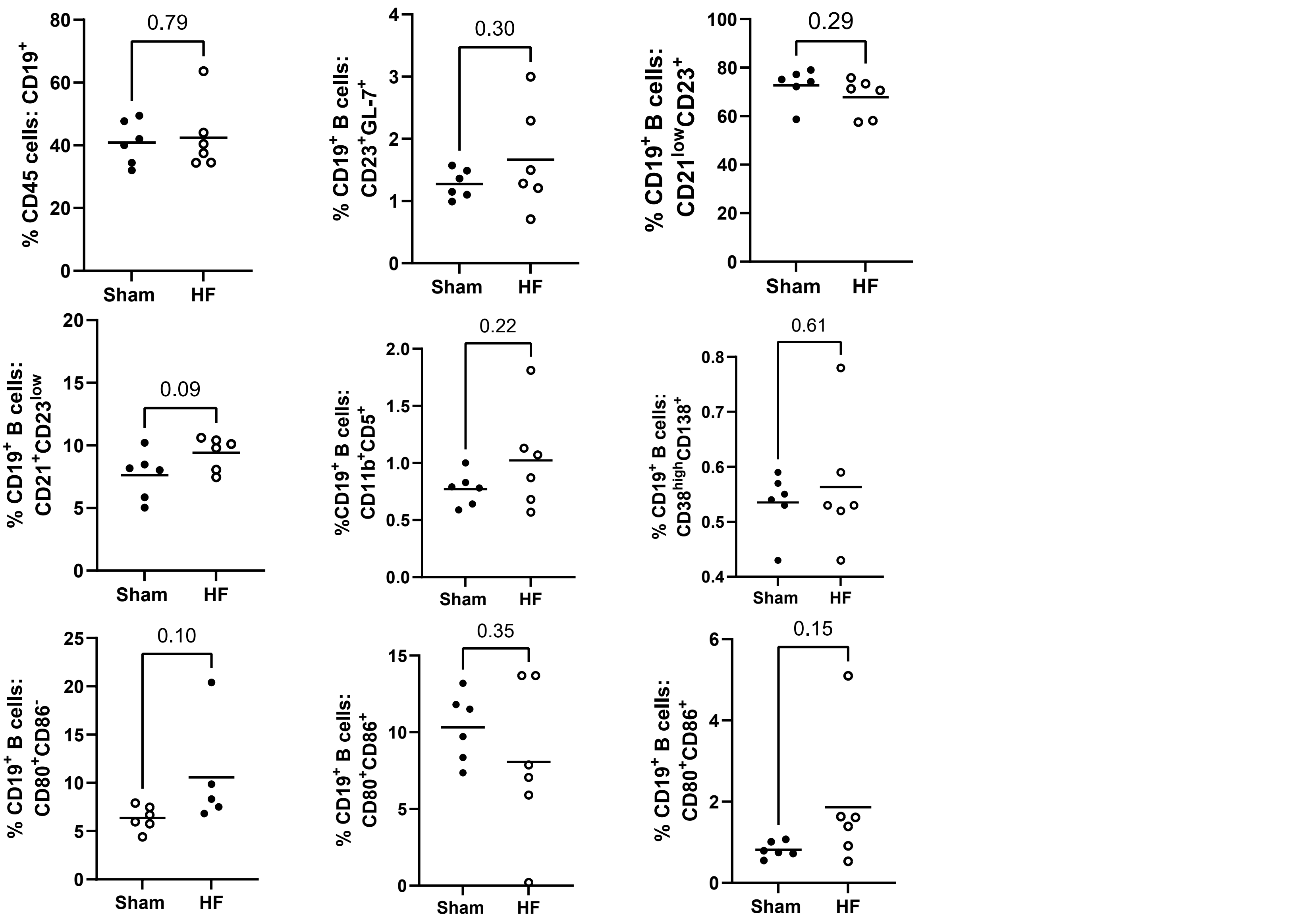


**Extended Data Fig.1: Adoptive transfer of splenic B cells did not induce changes in recipient splenic B cell subtypes.** Flow cytometry of the percentage of splenic B cell subtypes from naïve recipient mice eight-weeks following adoptive transfer from heart failure (HF) or sham-operated mice. Statistical analyses were performed using unpaired 2-tailed t-tests. n=6-7 male mice per group. Mean values are represented by horizontal lines. Marginal Zone B cells- CD21$^{+}$CD23$^{low}$; Follicular B cell- CD21$^{low}$CD23$^{+}$; Germinal Center B cells- CD23$^{+}$GL-7$^{+;}$ B1a B cells- CD11b$^{+}$ CD5$^{+}$; Plasma Cells - CD38high CD138$^{+}$; CD80 and CD86$^{+}$ were investigated as markers of B cells activation and both single positive for either marker and double positive cells are reported

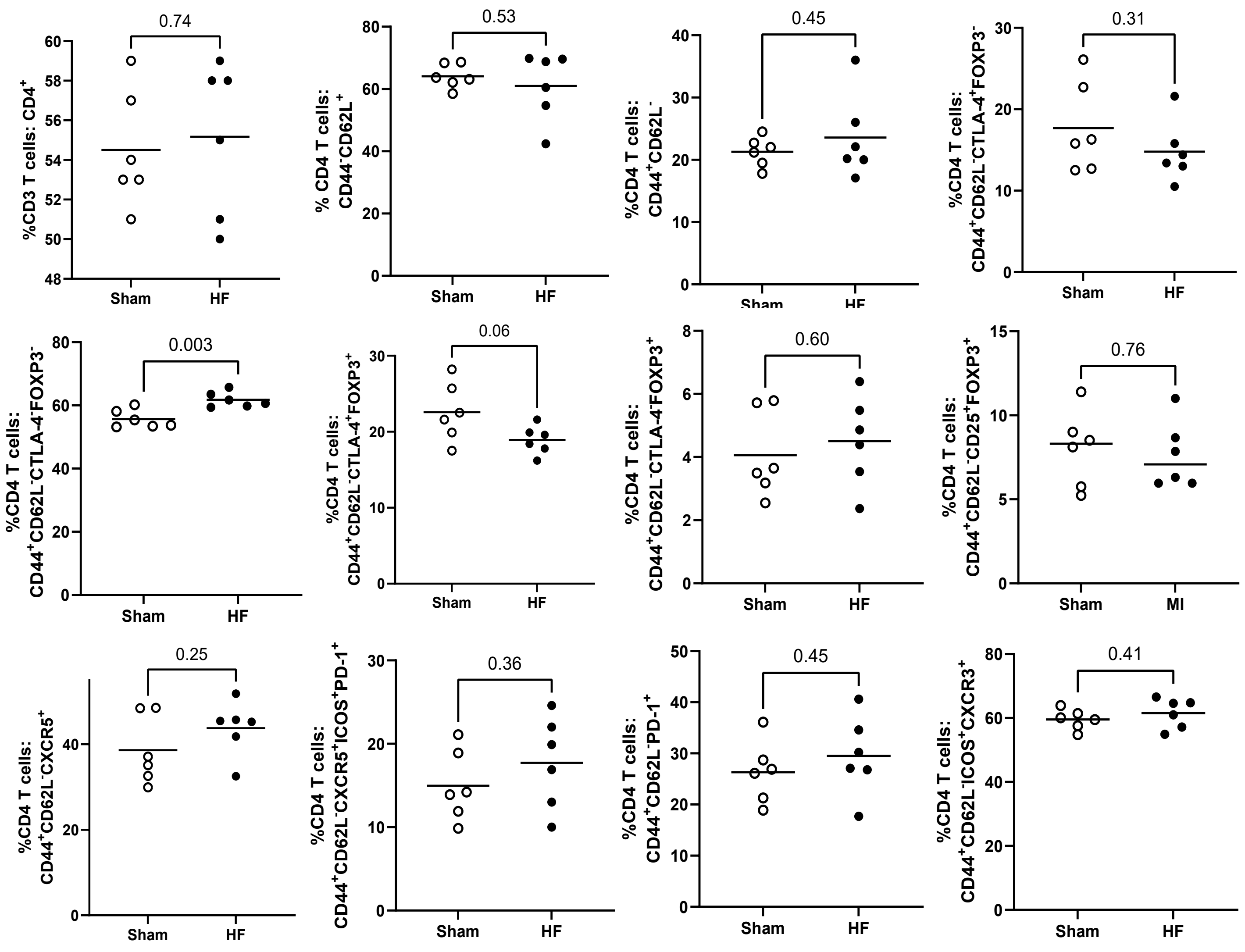


**Extended Data Fig.2: Adoptive transfer of splenic B cells induced minor changes in splenic CD4 T cell subtypes.** Flow cytometry of spleens from naïve recipient mice only showed a minor increase in the percentage of CTLA-4- FOXP3- memory CD4 T cells 8 weeks after adoptive transfer of splenic B cells from heart failure mice. Statistical analyses were performed using unpaired 2-tailed t-tests. n=6 male mice per group. Mean values are represented by horizontal lines

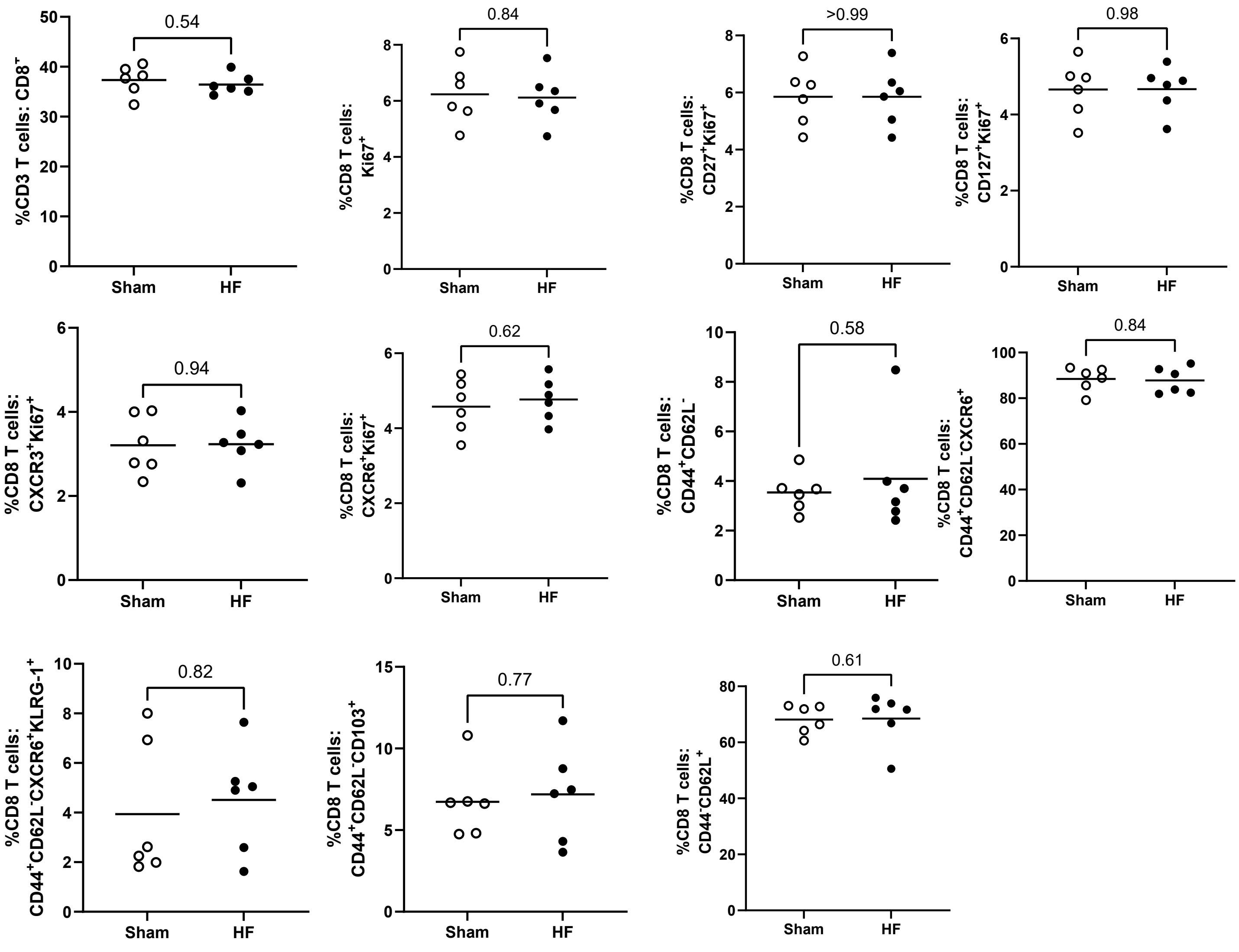


**Extended Data Fig.3: Adoptive transfer of splenic B cells did not induce changes in splenic CD8 T cell subtypes.** Flow cytometry of spleens from naïve recipient mice showed no changes in CD8 T cells 8 weeks after adoptive transfer of splenic B cells. Statistical analyses were performed using unpaired 2-tailed t-tests. n=6 per group. Mean values are represented by horizontal lines.

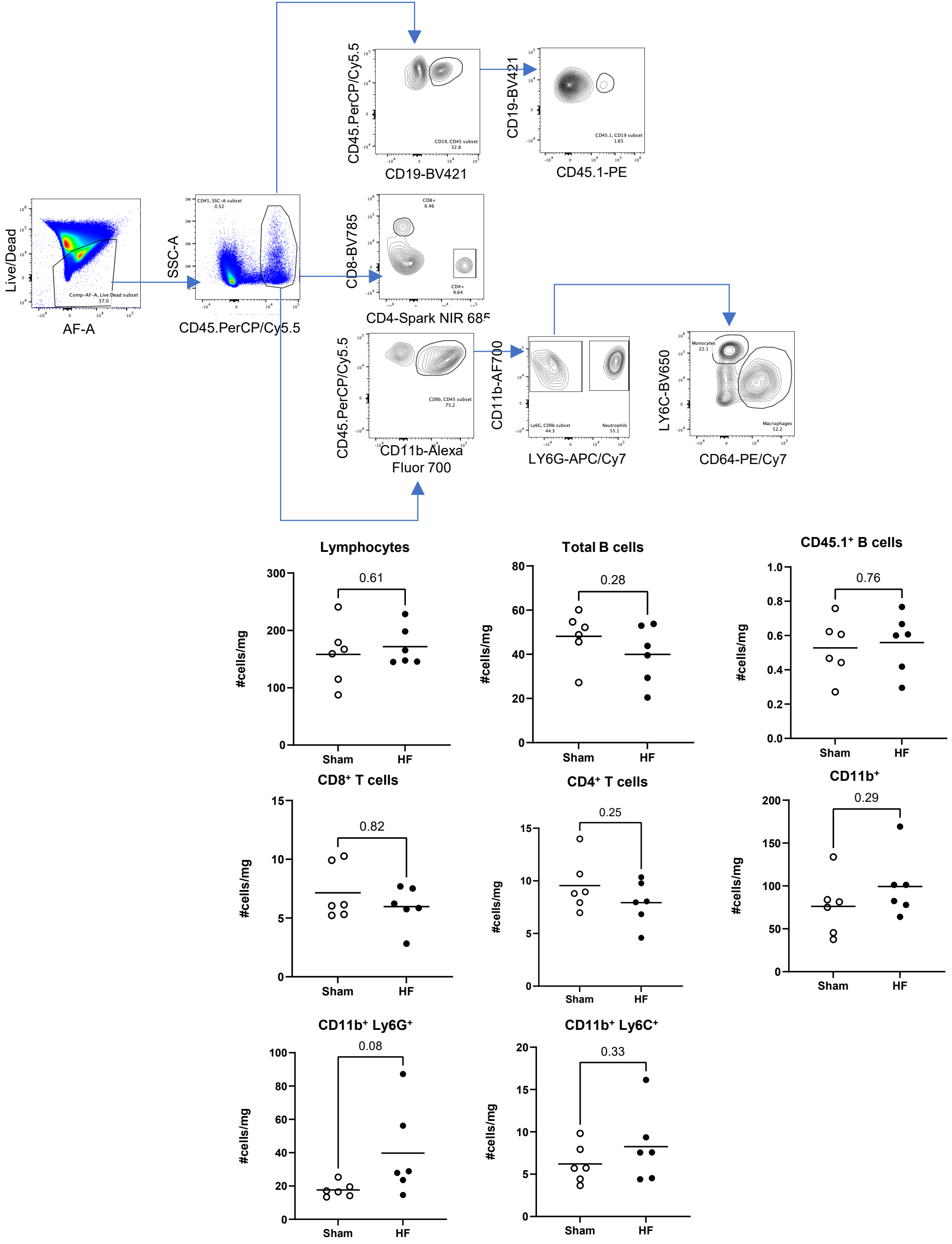


**Extended Data Fig.4: Adoptive transfer of splenic B cells from HF mice did not induce changes in <u>cardiac</u> immune cell populations of recipients.** a) Representative gating strategy b) Cardiac immune cell populations of recipient mice eight weeks post-adoptive transfer. Statistical analyses were performed with unpaired Mann-Whitney tests. n = 6 male mice per group. Mean values are represented by a horizontal mark.

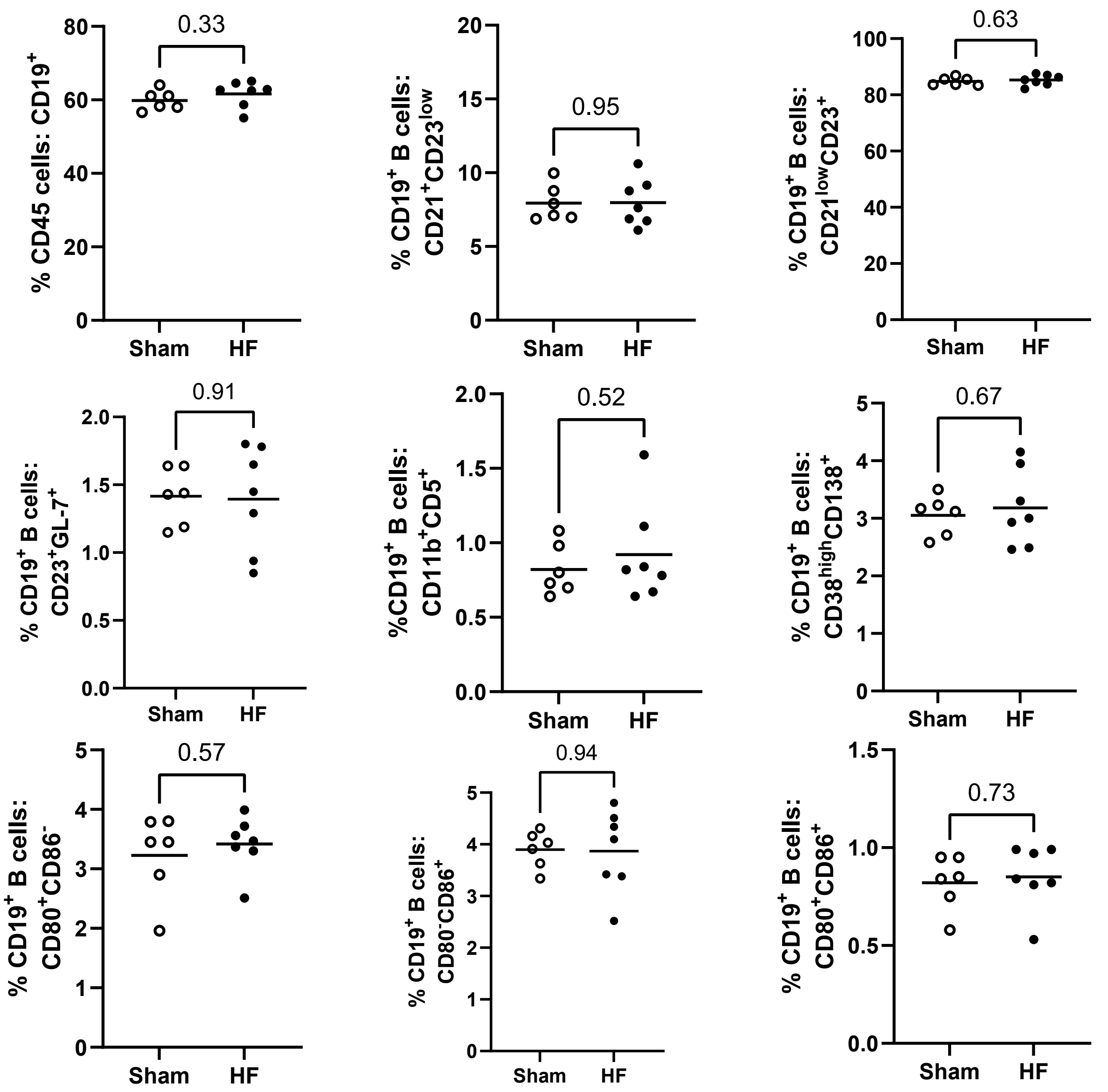


**Extended Data Fig.5: The prevalence of splenic B cell subtypes is unchanged in ischemic heart failure.** Flow cytometry of the percentage of splenic B cell subtypes from wildtype mice four weeks following permanent coronary artery ligation (MI) or sham surgery. Statistical analyses were performed using unpaired 2-tailed t-tests. n=6-7 male mice per group. Mean values are represented by horizontal lines. Marginal Zone B cells- $CD21^{+}CD23^{low}$; Follicular B cell- $CD21^{low}CD23^{+}$; Germinal Center B cells- $CD23^{+}GL\text{-}7^{+;}$ B1a B cells- $CD11b^{+}$ $CD5^{+}$; Plasma Cells - CD38high $CD138^{+}$; CD80 and $CD86^{+}$ were investigated as markers of B cells activation and both single positive for either marker and double positive cells are reported.
