## Supplementary Data for "Splenic follicular B cells promote adverse cardiac remodeling after myocardial infarction via MHC II-dependent antigen presentation"

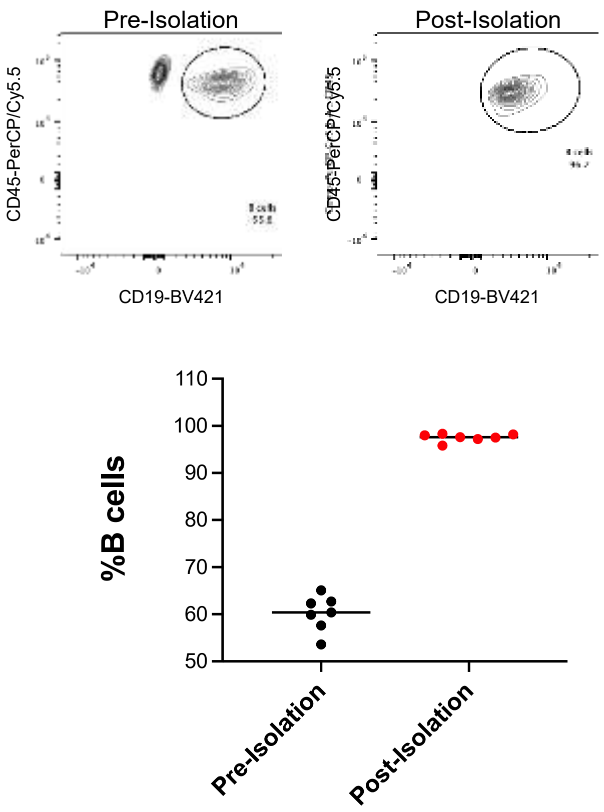


**Supplementary Fig.1: Representative B cell isolation for adoptive transfer studies.** Flow cytometry representative gating strategy showing CD19+ splenic B cells post isolation. Median value are presented

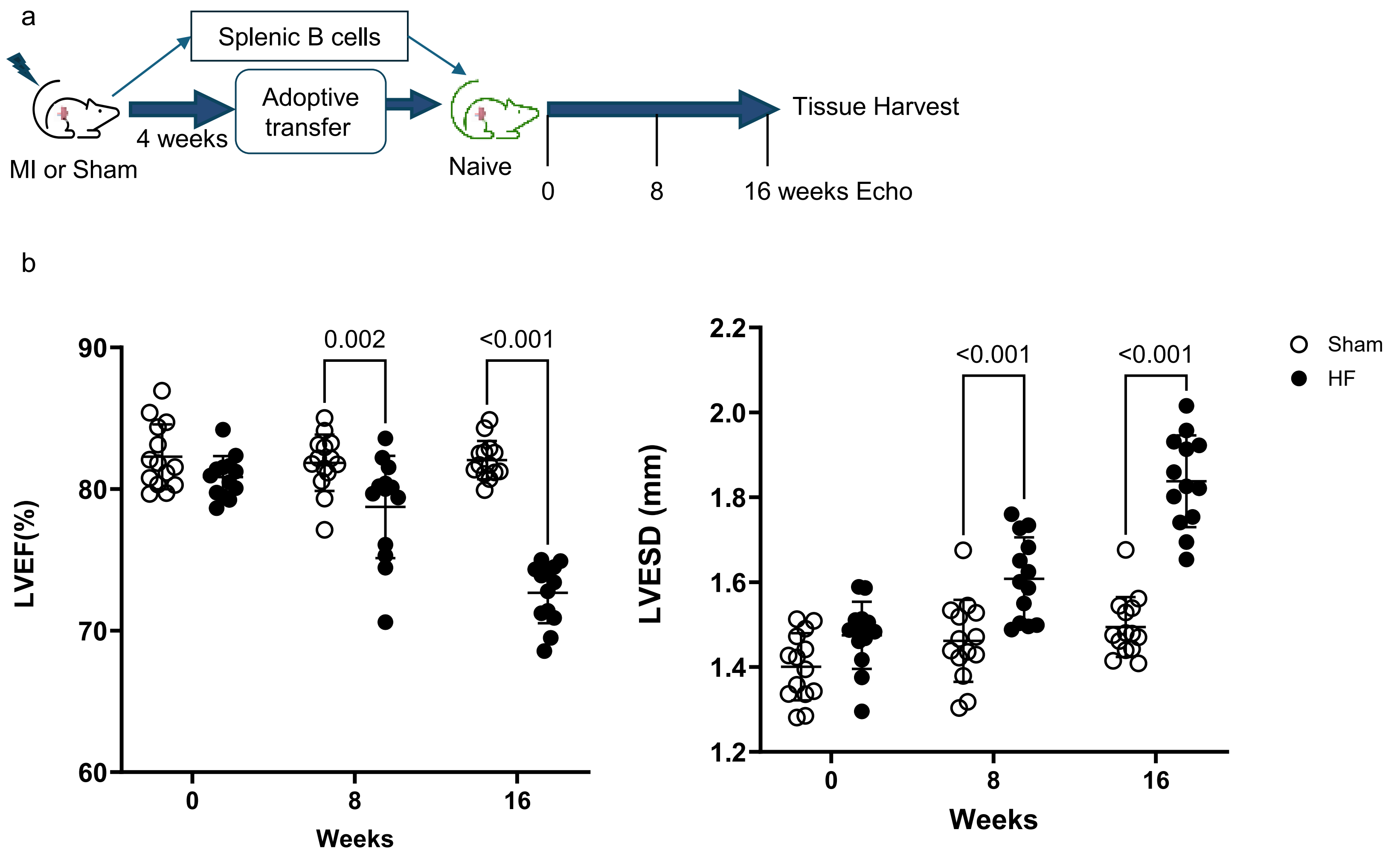


**Supplementary Fig.2: Adverse cardiac remodeling is sustained 16 weeks after adoptively transferred isolated heart failure splenic B cells** **a)** Schema for adoptive transfer studies. Four weeks following permanent coronary artery ligation or sham surgery, isolated splenic B cells were transferred to naïve recipient mice. Recipient mice underwent serial transthoracic echocardiography over 16 weeks followed by tissue harvest. **b)** On echocardiography, adoptive transfer of isolated splenic B cells from heart failure (HF) mice resulted in significant reduction in left ventricular ejection fraction (LVEF) and increase in left ventricular end-systolic diameter (LVESD) in naïve recipient mice, which was sustained at 16 weeks. Mean values ± SD are represented. Two-way ANOVA corrected with original FDR of Benjamini and Hochberg test.

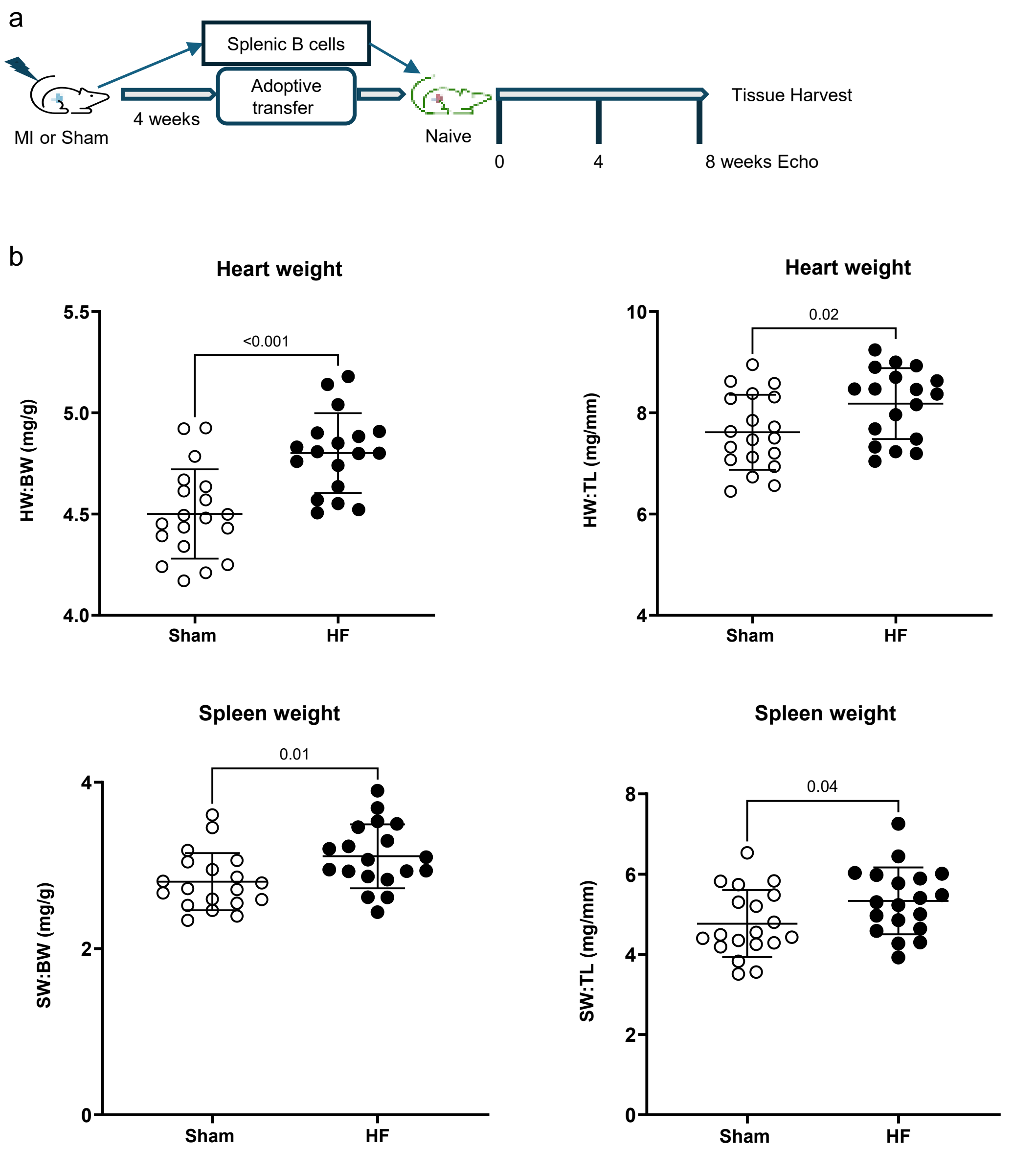


**Supplementary Fig.3: Combined analyses from independent splenic B cells adoptive transfer studies a)** Schema for adoptive transfer of isolated B cells. **b)** Combined gravimetric data from 3 independent experiments of recipient mice normalized to body weight and tibia length (n = 21 per group). Statistical analyses were performed with unpaired, 2-tailed t-tests. Mean values ± SD are represented. HW = heart weight. BW = body weight. SW = spleen weight. TL = tibia length.

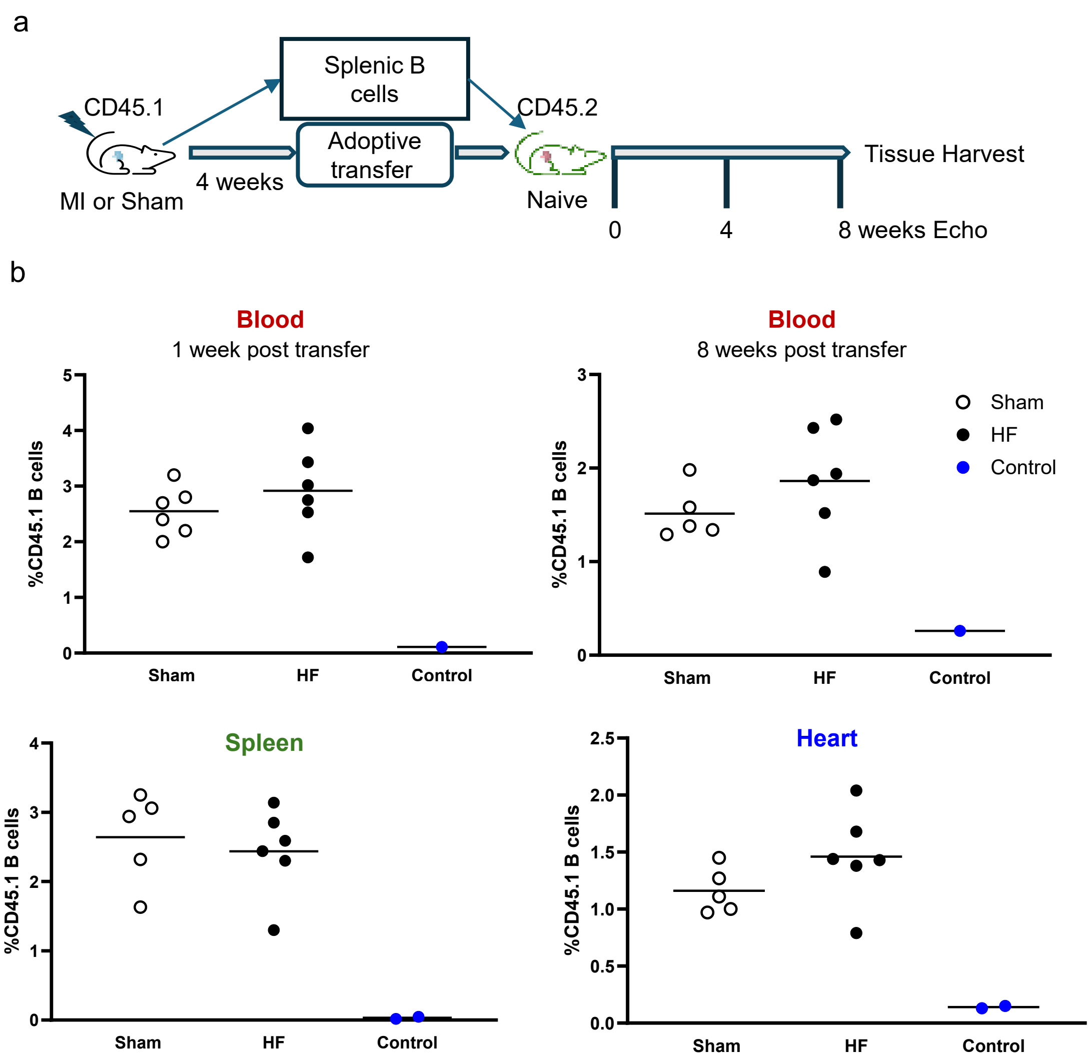


**Supplementary Fig.4: Validation of CD45.1 detection in analyzed samples a)** Schema for adoptive transfer of splenic B cells from CD45.1 mice post-Ischemic heart failure or sham to naïve CD45.2 mice. **b)** Flow cytometry with representative gating strategy showing presence of CD45.1 donor B cells in recipients', peripheral blood, spleen, and hearts. Non-recipient mice are included as an additional control. Horizontal bars represent mean values.

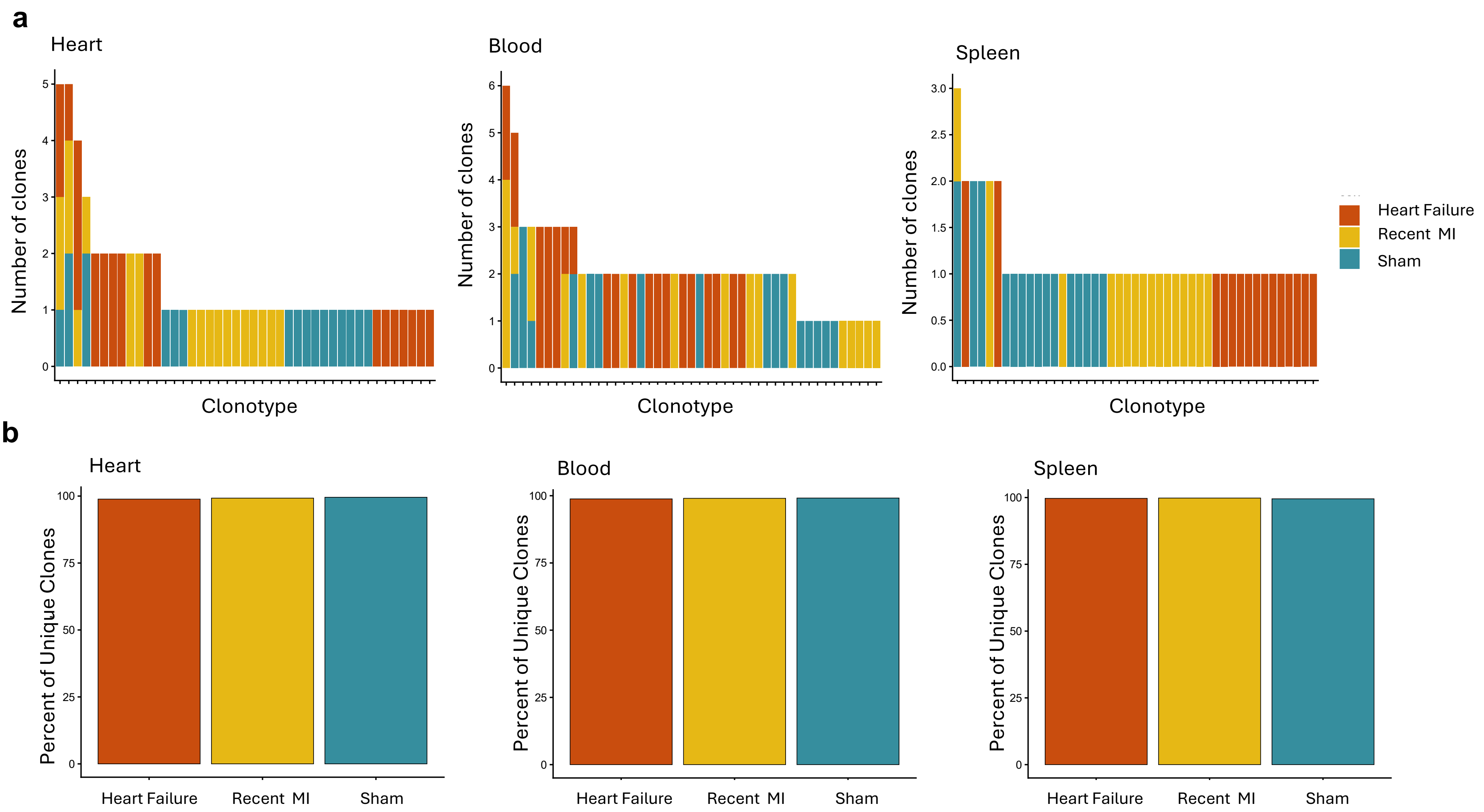


**Supplementary Fig.5: Myocardial infarction does not produce clear B cell clonal expansion a**) Stacked bar plot of the top 15 most abundant clonotypes in mice post recent MI (4 days post-MI), heart failure (4 weeks post-MI) or sham surgery, showing little to no clonal expansion and no difference in the distribution between heart failure and sham. **b**) Percent of unique clones showed nearly 100% of B cells in all conditions are unique, suggesting a lack of clonal expansion.

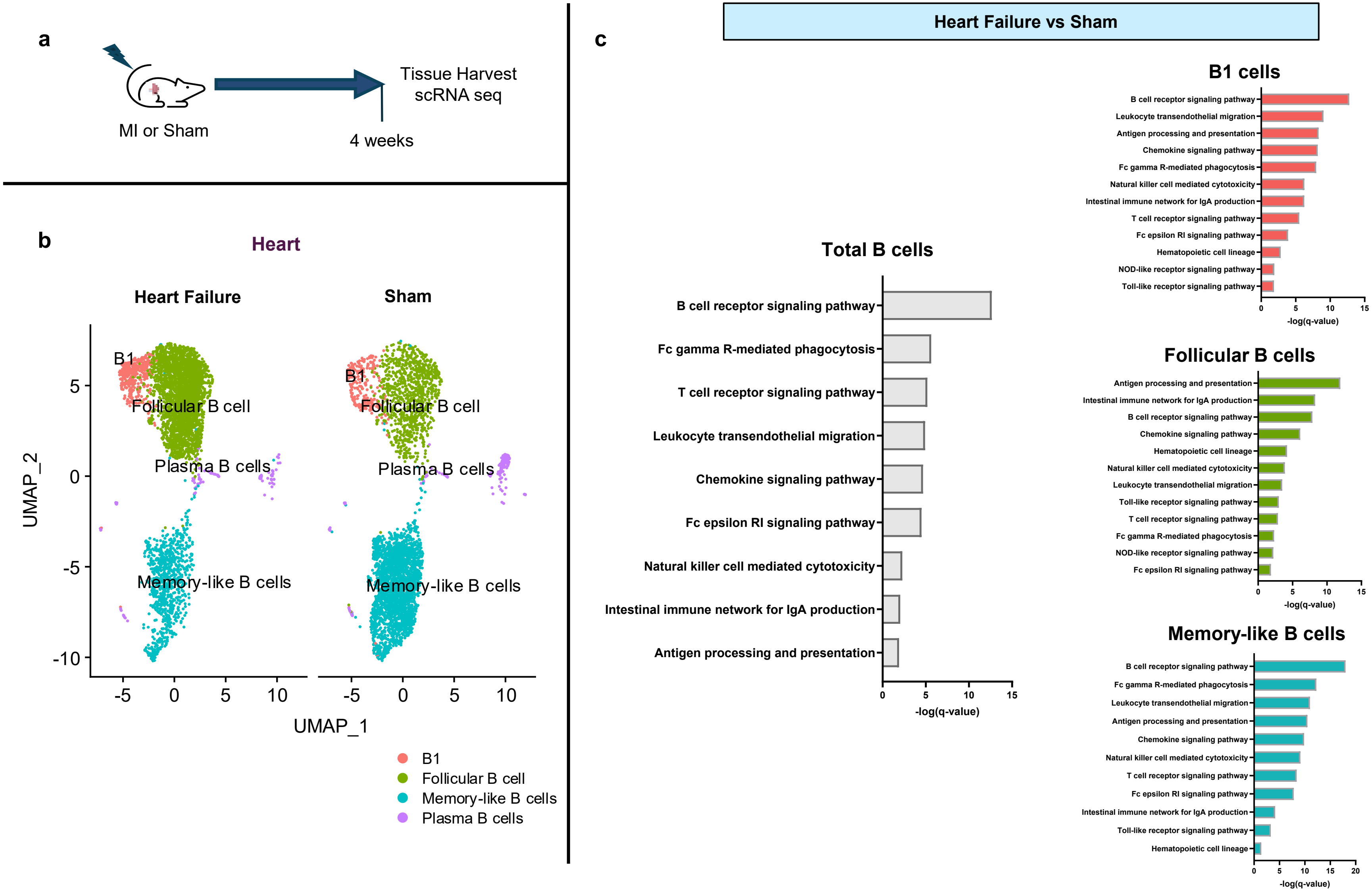


**Supplementary Fig.6: Ischemic heart failure results in dysregulation of antigen processing and presentation in cardiac B cells. a)** Schema for single cell sequencing of cardiac B cells from mice four weeks post-MI ("heart failure") or sham surgery. **b**) UMAP plots visualizing cardiac B cell sub-types. **c**) KEGG pathway analysis of top 500 differentially expressed genes in cardiac B cells from heart failure mice compared to sham mice (absolute fold-change $\leq$-1$\geq$1, $p<0.05$). Immune system pathways for selected B cell subtypes with q-value<0.05 are shown.

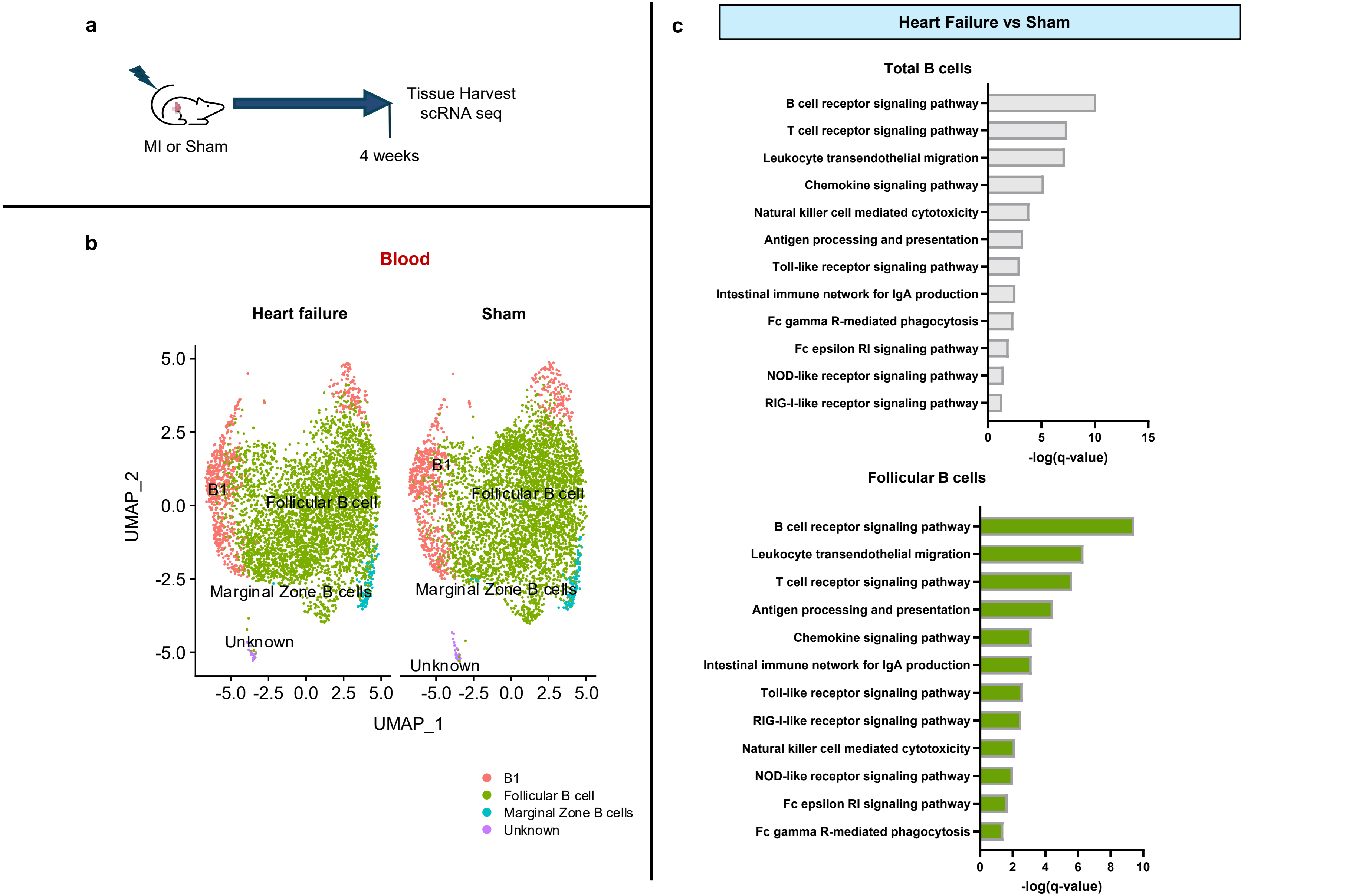


**Supplementary Fig.7: Ischemic heart failure results in chronic dysregulation of antigen processing and presentation in peripheral blood B cells. a**) Schema for single cell sequencing of peripheral blood B cells from mice four weeks ("heart failure") after permanent coronary ligation, or sham surgery. **b**) UMAP plots visualizing peripheral blood B cell sub-types. **c**) KEGG pathway analysis of differentially expressed genes in peripheral blood B cells from heart failure mice compared to sham mice (absolute fold-change ≤-1≥1, p<0.5). Immune system pathways with q-value<0.05 are shown.

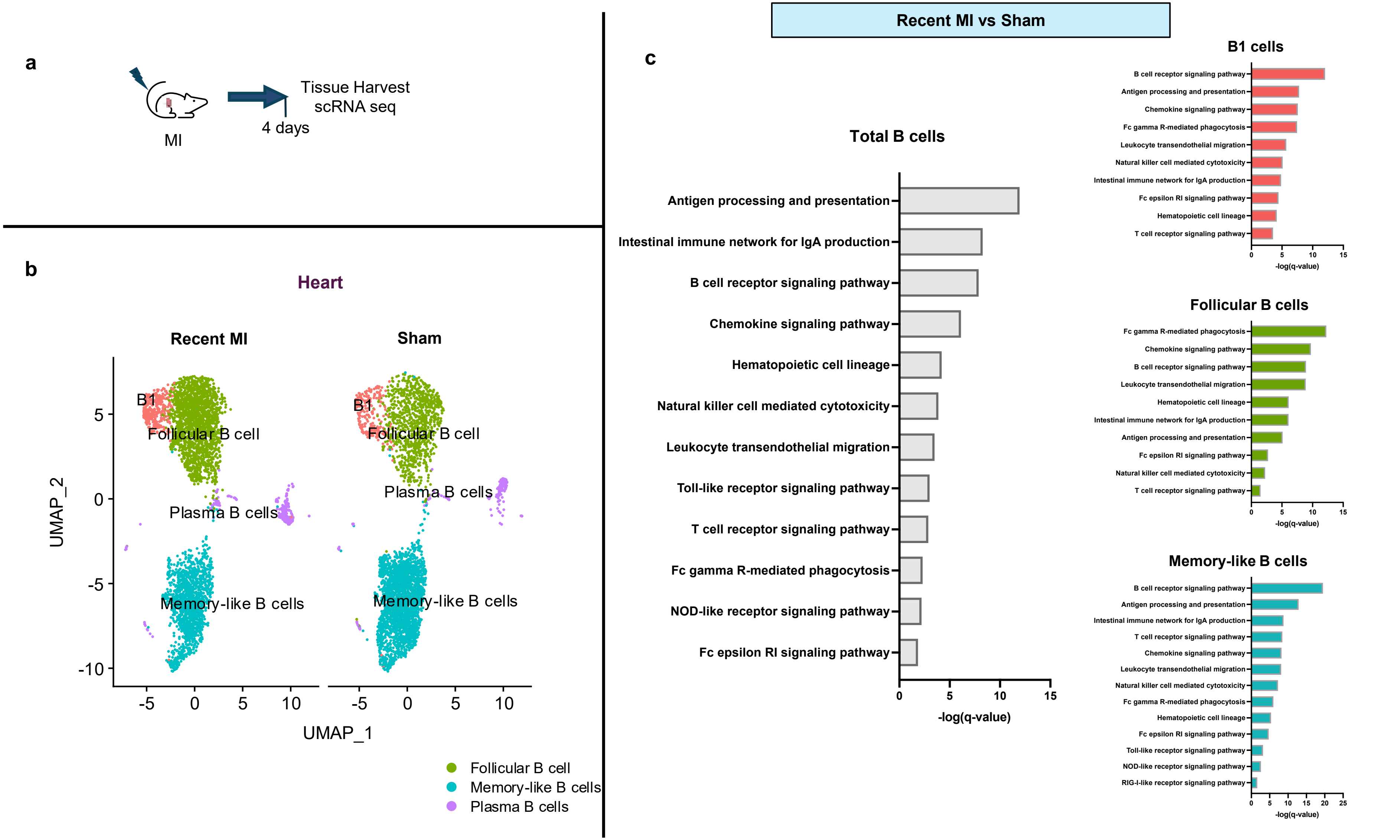


**Supplementary Fig.8: <u>Acute myocardial injury</u> results in dysregulation of antigen processing and presentation in <u>cardiac B cells</u>: analysis of recent MI vs sham a)** Schema for single cell sequencing of B cells from mice 4 days after MI ("recent MI"). B cells from mice 4 days after MI ("recent MI") were compared to B cells from mice 4 weeks after sham surgery. **b)** UMAP plots visualizing cardiac B cell sub-types. **c)** KEGG pathway analysis of differentially expressed genes in cardiac B cells from recent MI mice compared to sham mice (absolute fold-change $\leq -1 \geq 1$, $p<0.5$). Pathways with $FDR<0.05$ are shown.

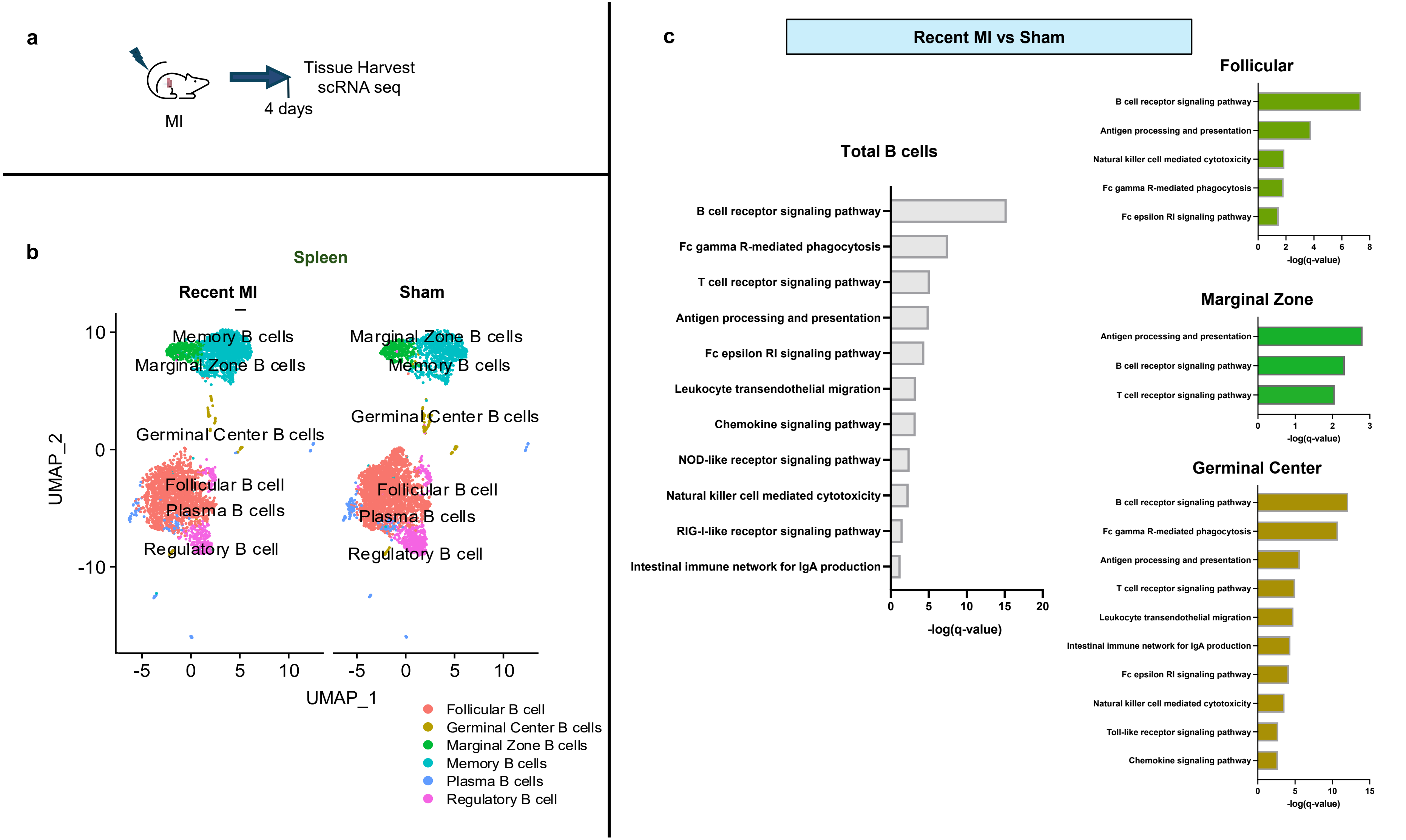


**Supplementary Fig.9: Acute myocardial injury results in dysregulation of antigen processing and presentation in splenic B cells. a)** Schema for single cell sequencing of B cells from mice 4 days after myocardial infarction ("recent MI") after permanent coronary ligation or 4 weeks after sham surgery. **b)** UMAP plots visualizing splenic B cell sub-types. **c**) KEGG pathway analysis of differentially expressed genes in splenic B cells from recent MI mice compared to sham mice (absolute fold-change $\leq$-1$\geq$1, p<0.5). Pathways with FDR<0.05 are shown.

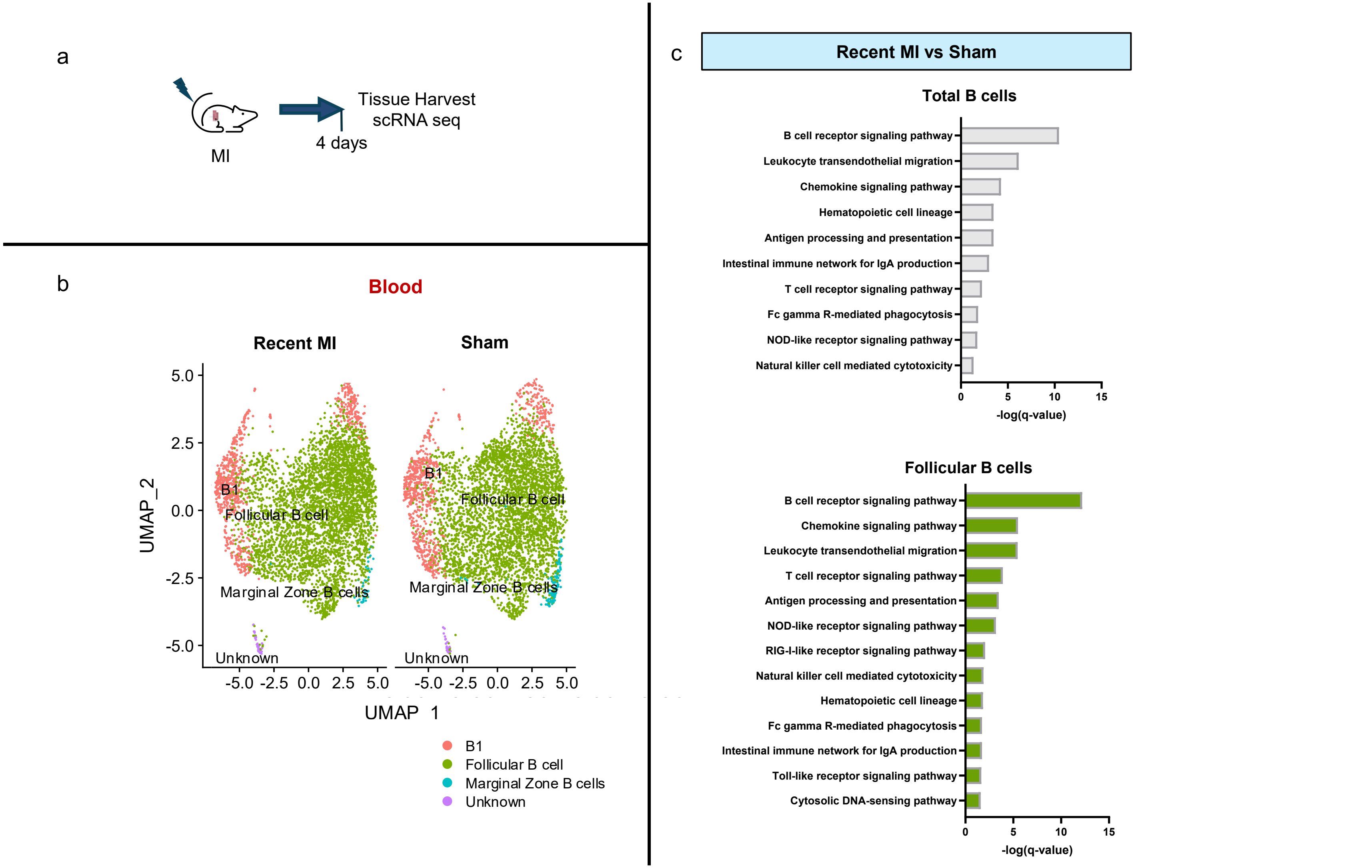


**Supplementary Fig.10: Acute myocardial injury results in dysregulation of antigen processing and presentation in peripheral blood B cells** **a**) Schema for single cell sequencing of B cells from mice 4 days ("recent MI") or 4 weeks ("heart failure") after permanent coronary ligation, or sham surgery. **b**) UMAP plots visualizing peripheral blood B cell sub-types. **c**) KEGG pathway analysis of differentially expressed genes in cardiac B cells from HF mice or recent MI mice compared to sham mice (absolute fold-change ≤-1≥1, p<0.5). Pathways with FDR<0.05 are shown.

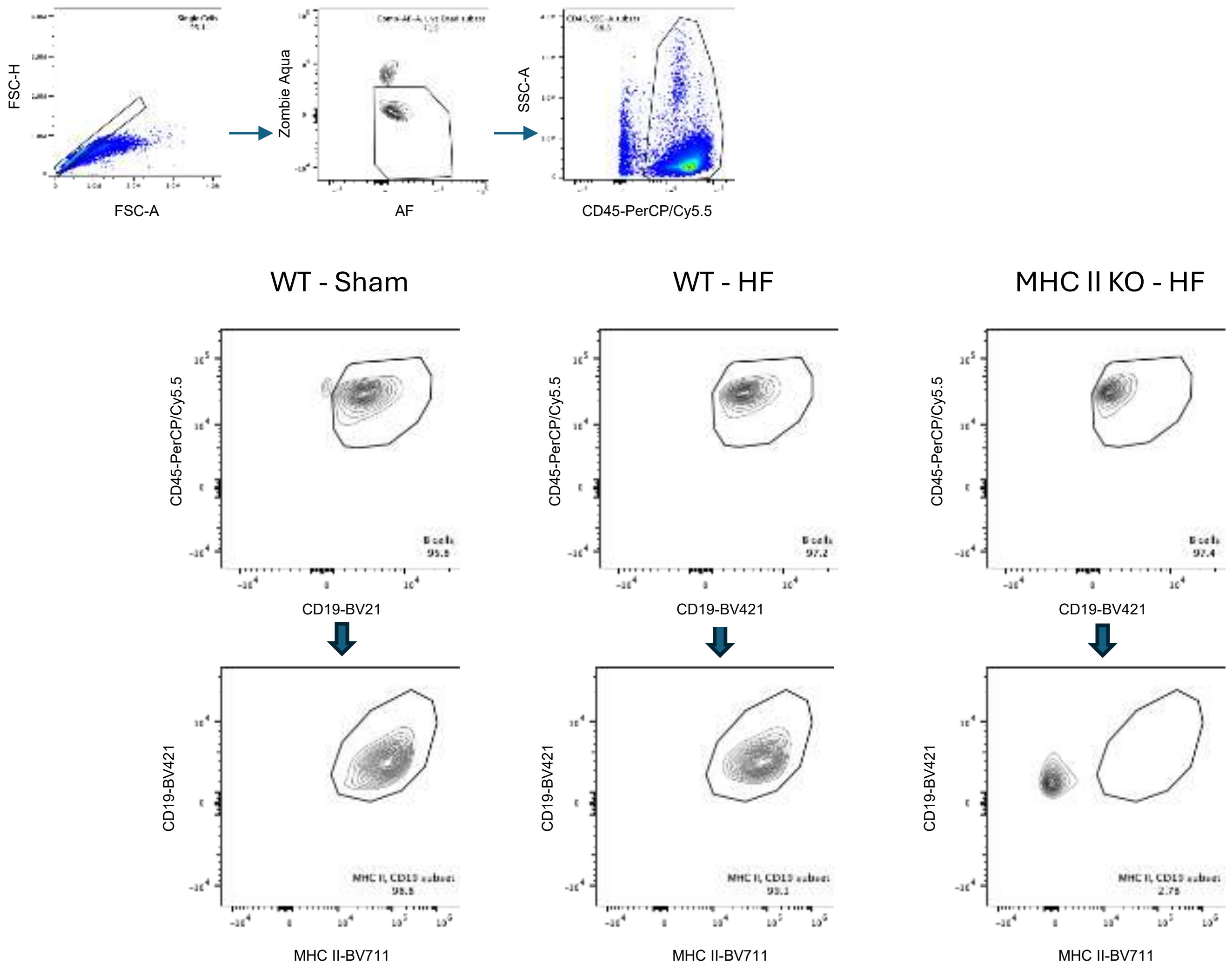


**Supplementary Fig.11: Confirmation of MHC class II deletion on splenic B cells in transgenic mouse model.** Splenic B cell isolation from *Cd19*$^{tm1(cre)Cgn/-}$*H2-Ab1*$^{b\text{-tm1Koni}/\,b\text{-tm1Koni}}$ mice and wildtype mice prior to adoptive transfer.

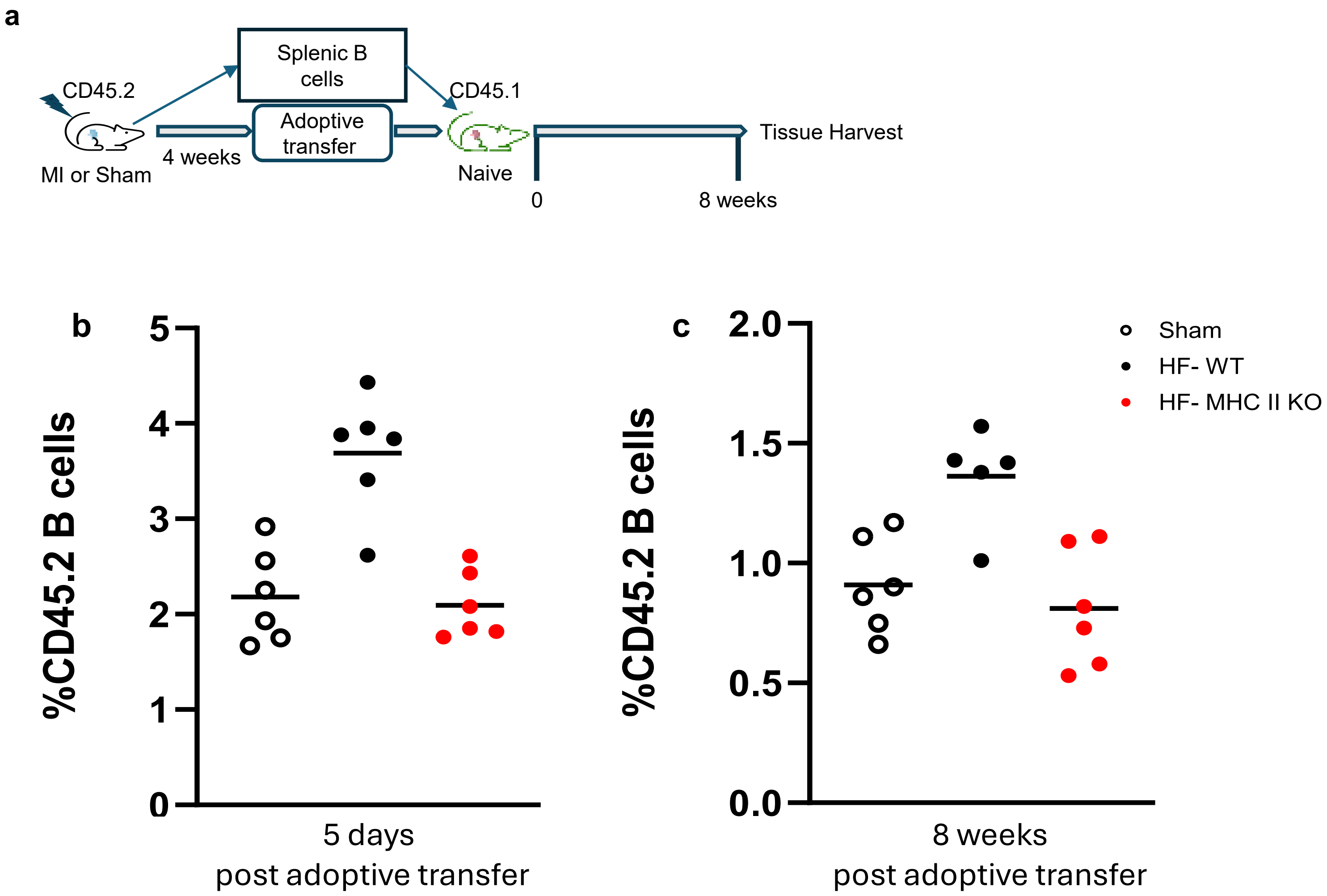


**Supplementary Fig.12: Confirmation of presence of donor B cells from MHC II-deficient mice in naïve recipient wildtype mice. a**) Schema for adoptive transfer of isolated B cells. Quantification of donor-derived CD45.2 B cells in recipients' peripheral blood **b**) 5 days and **c**) 8 weeks post-adoptive transfer. Mean values are represented.

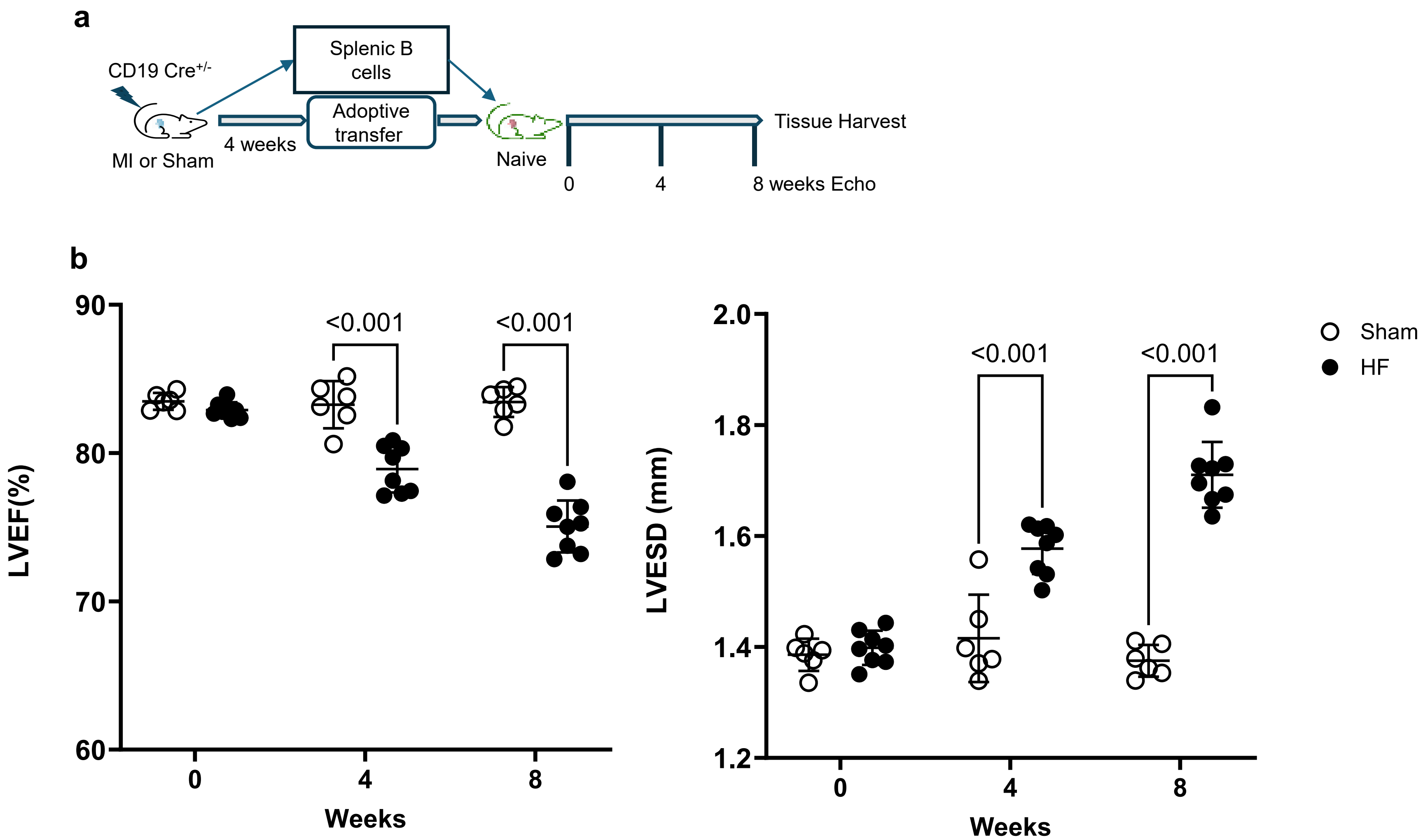


**Supplementary Fig.13: CD19 Cre Donors a**) Schema for adoptive transfer of isolated B cells. **b**) Serial echocardiographic data of recipient mice over eight-week period post-adoptive transfer of isolated splenic B cells (n = 6-8 per group). Statistical analyses were performed with Two-way ANOVA and Mixed-effects analysis corrected with original FDR of Benjamini and Hochberg test. Mean values ± SD are represented.

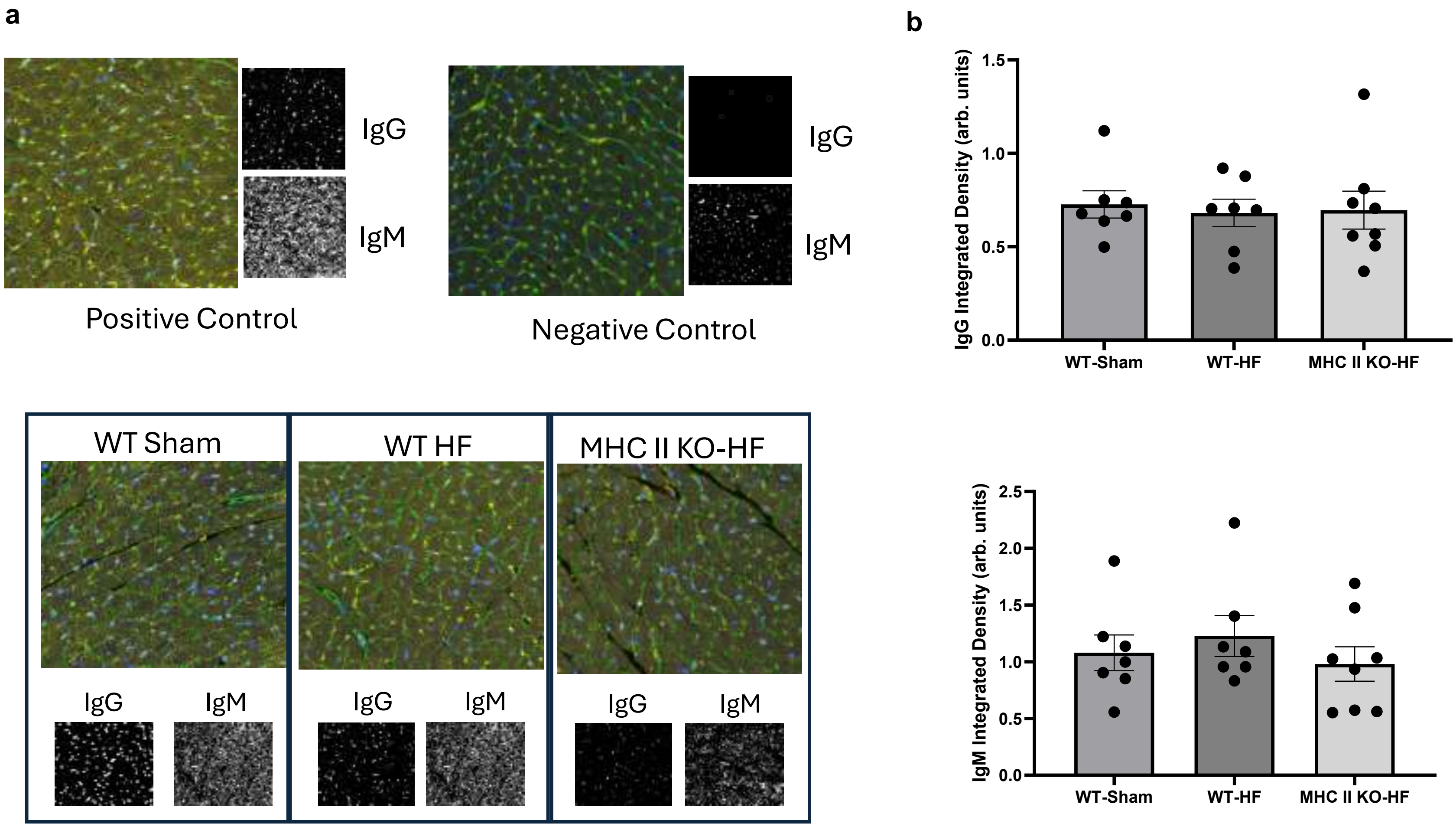


**Supplementary Fig.14: Adoptive transfer of HF splenic B cells does not increase cardiac autoantibodies in recipients a**) Representative images of □MT heart sections incubated with serum from recipients of WT sham, WT MI and MHC II KO MI splenic B cells, and stained with Anti-IgG and IgM antibodies. Positive control was serum from old (70 weeks old mice). **b**) Quantified data shows no difference between groups in IgG and IgM autoreactivity, n=4 mice per group; 2 technical replicates. Data presented as Mean ± SEM. One-way ANOVA was used to determine statistical significance.

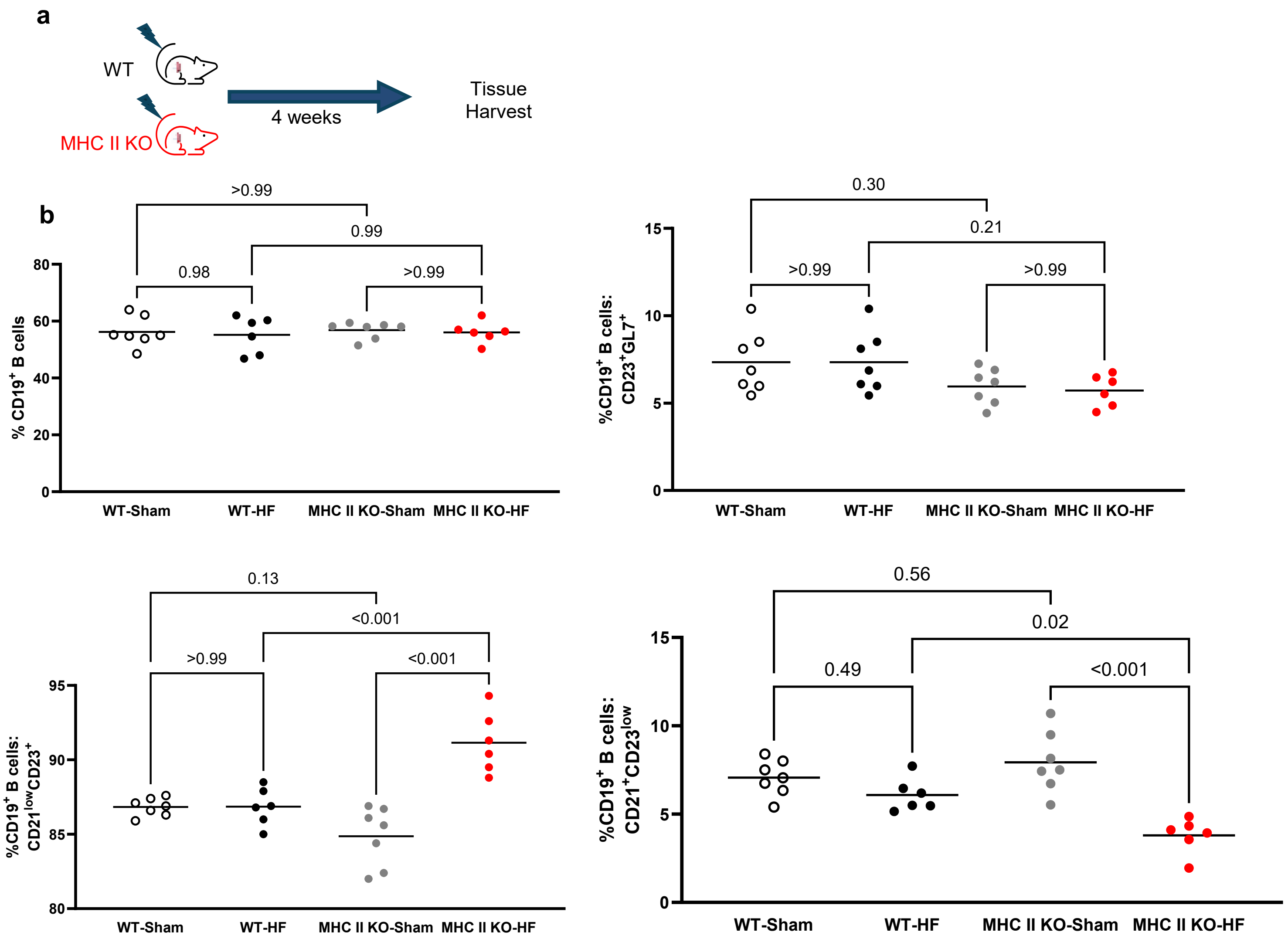


**Supplementary Fig.15: B cell specific MHC-II deletion is associated with follicular splenic B cells expansion after ischemic myocardial injury a**) Schema for timing of harvesting of splenic B cells from *Cd19*$^{tm1(cre)Cgn/-}$*-H2-Ab1*$^{b\text{-tm1Koni}/\ b\text{-tm1Koni}}$ (MHC II KO) and wildtype (WT) mice following permanent coronary artery ligation or sham surgery. **b**) Splenic follicular B cells are increased in MHC II KO mice and splenic marginal zone B cells are decreased post ischemic myocardial injury. Statistical analyses were performed using unpaired 2-tailed t-tests. n=6-7 male mice per group. Mean values are represented. Follicular B cell- CD21$^{low}$CD23$^{+}$; Marginal Zone B cells- CD21$^{+}$CD23$^{low}$; Germinal Center B cells- CD23$^{+}$GL-7$^{+}$

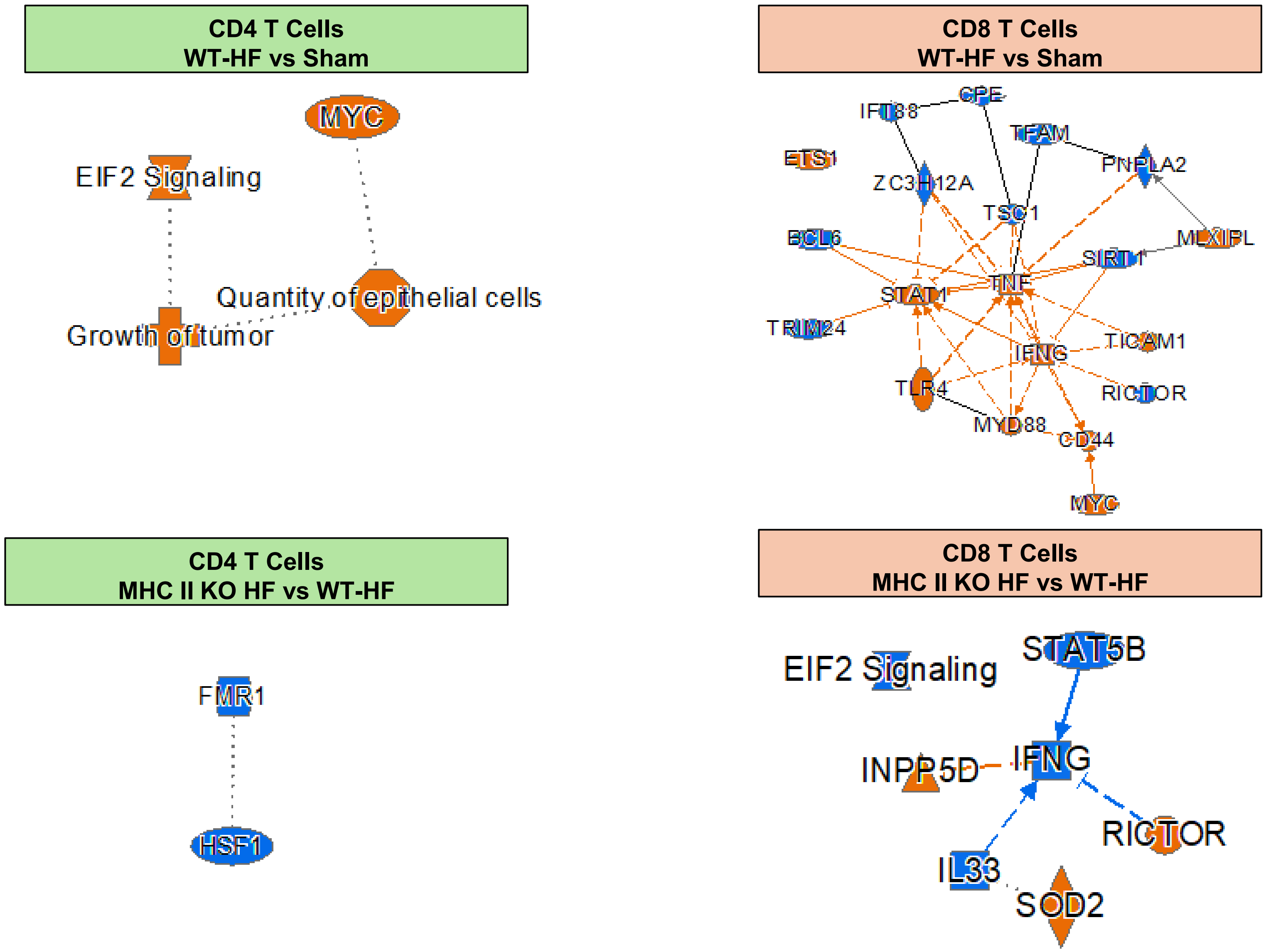


**Supplementary Fig.16: Signaling network analysis of chronic gene expression changes in myocardial T cells produced by adoptive transfer of splenic B cells from mice with ischemic HF, in the presence or absence of MHC-II on B cells** Ingenuity Pathway Analysis (IPA) based analysis of upstream regulators and signaling cascades. Splenic B cells activate IFN gamma related pathways in myocardial CD8 T cells and HSF1 related pathways in CD4 T cells. Deletion of MHC-II on transferred B cells prevents these effects and results in relative downregulation. Orange = upregulation, blue = downregulation
